# A knowledge graph-guided framework for multitask single-cell analysis

**DOI:** 10.64898/2026.04.25.720773

**Authors:** Bernadette Mathew, Akash Maity, Ritwik Ganguly, Abhishek Halder, Justin Thomas Sabu, Neha Gupta, Sreeram Chandra Murthy Peela, Swarnava Samantha, Stuti Kumari, Nilabja Bhattacharjee, Shalaka Joshi, Sudeep Gupta, Gaurav Ahuja, Mohammed Farooq, Ralf Herwig, Martin Vingron, Angshul Majumdar, Syed Hasan, Debarka Sengupta

## Abstract

Connecting cell identity to biological function is central to understanding how tumors change during treatment. Here, we introduce scKNIFE, a knowledge-guided nonnegative matrix factorization framework that integrates gene expression with a heterogeneous graph of functional annotations, metabolic reactions and metabolites, and signaling interactions. scKNIFE jointly learns cellular representations and biological-entity loadings, linking the factors that organize cells to their constituent genes and connected functions. Across breast-cancer cohorts, the learned representations retained broad cellular organization without using per-cell identity labels as factorization targets. scKNIFE identified baseline tumor programs associated with subsequent clinical response or T-cell clonotype expansion and resolved treatment-associated functional variation within immune and stromal populations. In a seven-patient triple-negative breast cancer atlas, analysis alongside tissue metabolomics identified distinct trajectories of inferred fatty acid activation and carnitine-associated activities, including higher myeloid L-octanoylcarnitine scores at biopsy in all seven patients. Communication analysis further resolved an endothelial-perivascular PDGF connection with lower inferred scores at surgery in six patients. scKNIFE brings cellular identity, metabolic programs and candidate intercellular communication into one representation, enabling functional interpretation of tumor heterogeneity across patients and treatment states.

---

Single-cell RNA sequencing (scRNA-seq) can resolve the cellular heterogeneity of tumors. In triple-negative breast cancer (TNBC), epithelial, stromal, and immune populations vary among patients and change during treatment ^1–3^. Treatment can also select or reprogram cellular states associated with resistance and recurrence ^4^. Interpreting these changes requires methods that retain cell-level variation and connect it to biological function and clinical context.

Most single-cell workflows analyze expression programs, metabolism, and cell–cell communication separately. Gene-set and latent-factor methods, including Spectra, expiMap, and Slalom, organize expression into interpretable programs ^5–7^. Metabolic methods infer reaction- or task-level activity, whereas communication methods combine ligand–receptor expression with curated interaction resources ^8,9^. Because these analyses produce separate outputs, relationships among cell identity, metabolism, signaling, and other biological processes must be assembled after model fitting.

A heterogeneous biological knowledge graph can encode these relationships before fitting by linking genes to reactions, metabolites, pathways, ontology terms, cell-state annotations, and signaling entities. Graph regularization can then let expression-supported nodes inform connected entities in the learned representation. This propagation is a model-derived inference, not direct evidence of causal regulation, physical interaction, or metabolic flux.

Non-negative matrix factorization (NMF) represents each cell as an additive combination of partially overlapping programs ^10,11^. Graph-regularized NMF encourages connected entities to have related latent loadings ^12^. Applying this approach to a heterogeneous graph requires mapping measured genes to graph nodes and distinguishing node loadings from cell-level factor use. It also requires separating expression-linked activity from scores inferred through graph connections.

We developed scKNIFE (knowledge graph-based NMF for inference of functional entities) to learn cellular representations and graph-linked biological activities within a common factorization. The model approximates gene expression as **X** *≈* **R**_*g*_**WH**, where **R**_*g*_ maps measured genes to graph nodes, **W** contains loadings for genes and functional entities, and **H** describes factor use across cells. Expression reconstruction anchors the model to measured transcription, while graph regularization links the loadings of connected entities. The product **WH** provides cell-resolved scores for genes, reactions, metabolites, pathways and signaling entities within the same latent representation. This construction allows a cellular program to be examined through both its expression-supported components and its connections to broader biological functions. The graph incorporates curated resources for metabolism, signaling, functional annotations and cellular identity ^9,13,14^.

We evaluated scKNIFE in four external breast-cancer datasets and the Tahoe drug-perturbation compendium. These analyses compared its representation and activity scores with established methods and examined treatment-associated metabolic patterns. We then analyzed 15 single-cell specimens from seven patients with TNBC, alongside paired biopsy and surgery tissue metabolomics. This longitudinal atlas let us examine epithelial programs, compartment-specific metabolic activity, one patient’s recurrence specimen, and candidate ligand–receptor communication.

## Results

### scKNIFE links cellular variation to biological programs through knowledge-guided factorization

scKNIFE defines latent factors over a heterogeneous biological graph, allowing measured genes and connected functional entities to share a common representation (Figure 1a). Each factor pairs a graph-node loading profile in **W** with a cell-level activity profile in **H**. These matched profiles connect the factors organizing cells to their constituent genes, reactions, metabolites and signaling entities, while **WH** provides the corresponding node activities in individual cells.

**Figure 1.**
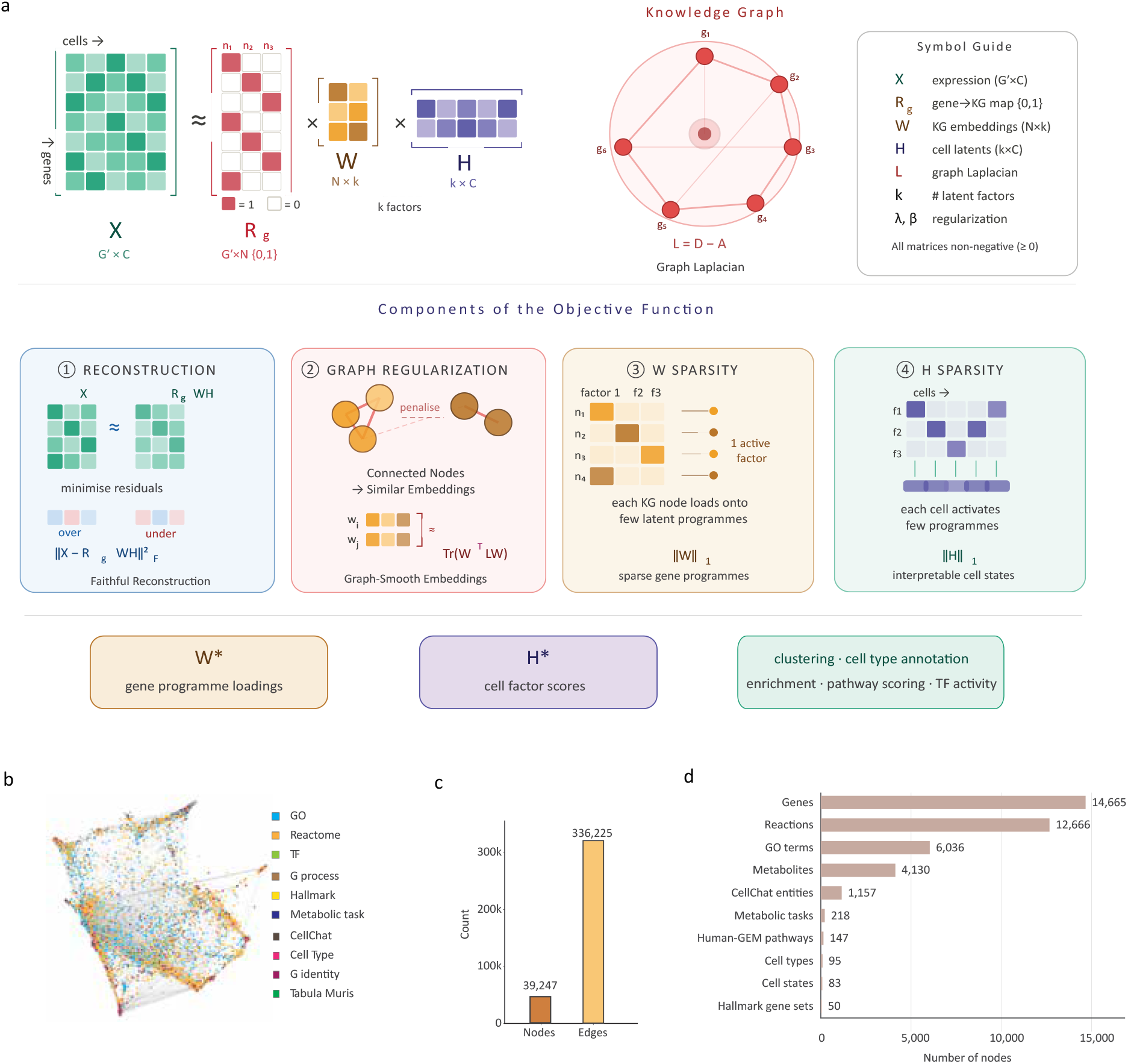
Knowledge-guided factorization and the scKNIFE graph. (a) Overview of the scKNIFE model. The gene-by-cell expression matrix X is approximated by RgWH, where Rg maps measured genes to graph nodes, W contains graph-node loadings on latent factors, and H contains factor usages across cells. Graph regularization uses the Laplacian *L* = *D − A* to encourage connected nodes to have similar loadings. Nonnegative W and H yield node activities through WH. The implementation combines expression reconstruction, graph smoothness and *L*_1_ penalties on W and H. (b) Schematic organization of the heterogeneous knowledge graph, integrating genes, metabolic reactions and tasks, metabolites, biological annotations and signaling entities. Colors distinguish knowledge sources or annotation categories. (c) Size of the expanded graph used for the in-house analyses: 39,247 nodes and 336,225 edges. (d) Node composition: 14,665 genes, 12,666 reactions, 6,036 GO terms, 4,130 metabolites, 1,157 CellChat entities, 218 metabolic tasks, 147 Human-GEM pathways, 95 cell-type annotations, 83 cell-state annotations and 50 Hallmark gene sets. Panels c,d describe the expanded in-house graph; the external benchmark graph contains 22,951 nodes and 216,079 edges. Curated cell-identity nodes encode prior knowledge, whereas observed per-cell identity labels are not factorization targets.Graph entities and fitted factors describe the model architecture and knowledge coverage. Panels c,d report graph counts.

The model approximates the gene-expression matrix as *X ≈ R*_*g*_*WH*: *R*_*g*_ maps measured genes to graph nodes, *W* contains node loadings across factors, and *H* describes factor use in each cell (Figure 1a). Expression reconstruction anchors the factors to observed transcriptional variation. Graph regularization encourages connected entities to share similar loadings, while sparsity penalties limit the contributions to each node and cell. Cells can use several factors, and each factor can include entities from multiple biological annotations. Factorization is anchored in gene-expression reconstruction, while curated cell-identity and cell-state information enters through the graph as biological context.

The knowledge graph connects genes to functional entities spanning biological processes, signaling interactions, metabolic reactions and tasks, and curated cell-type and cell-state annotations (Figure 1b). For the external benchmarks, the graph contained 22,951 nodes and 216,079 edges. We expanded this graph for the analyses of the TNBC study cohort by incorporating Human-GEM, the consensus human metabolic reconstruction ^13^, including explicit metabolite and metabolic-pathway nodes. The expanded graph contained 39,247 nodes and 336,225 edges (Figure 1c; Supplemental Table 2). Its metabolic component comprised 12,666 reactions, 4,130 metabolites, 218 metabolic tasks and 147 Human-GEM pathways, alongside 14,665 genes, 6,036 Gene Ontology terms, 1,157 CellChat interaction entities ^9^, 95 cell-type annotations, 83 cell-state annotations and 50 Hallmark gene sets (Figure 1d). These connections allow gene-expression information to inform the representation of linked biological processes at several levels of organization.

The model provides complementary views of cellular variation. *H* gives each cell’s factor usage and can be used to build neighborhoods and clusters; *W* links each factor to genes and other graph entities (Figure 1a). Because expression and graph structure shape the factors together, a factor can connect overlapping annotations, including cell identity, signaling and metabolism. The product *WH* gives an inferred activity score for each graph node in each cell. For non-gene entities, these scores summarize transcriptomic support across modeled relationships and provide inferred activity estimates that can be interpreted alongside direct measurements of metabolite abundance or flux.

We next assessed whether the graph-guided representation retained cellular organization and supported biological interpretation across independent datasets.

### scKNIFE retains cellular organization across cohorts while representing broad biological knowledge

We first characterized the breadth of biological knowledge represented in scKNIFE and the comparator configurations. The benchmark graph encoded 7,521 configured biological-program category memberships; the expanded in-house graph encoded 7,668 after adding 147 Human-GEM pathways (Figure 2b). The two graphs contained 22,951 and 39,247 nodes, respectively (Supplemental Table 2). The tested comparator configurations used 184 gene sets or factors for SPECTRA, 843 gene sets for expiMap, and 118 gene sets or factors for Slalom. These counts describe implementation inputs, not equivalent model units across methods ^5–7^.

**Figure 2.**
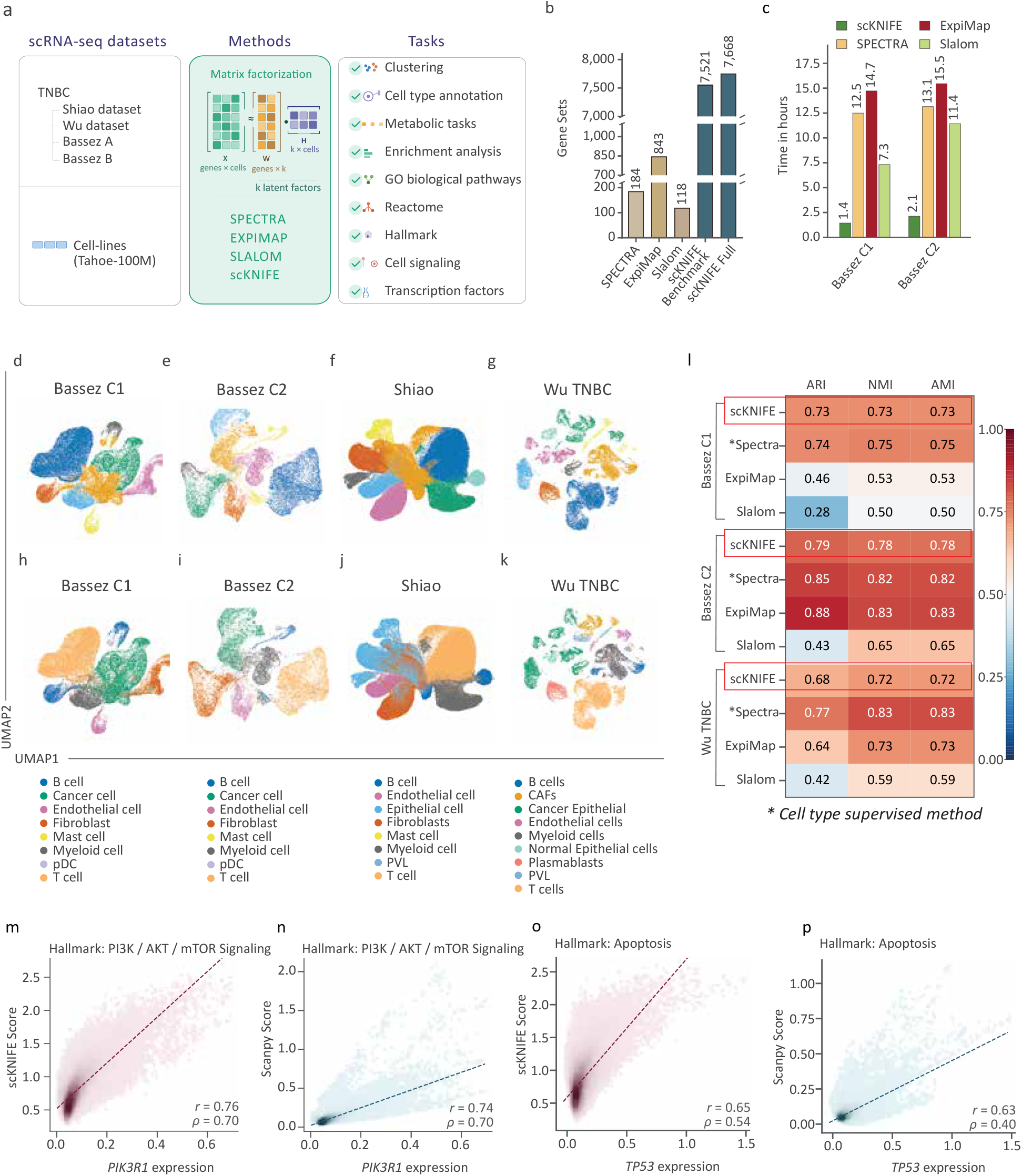
Broad knowledge-guided scKNIFE representations preserve cellular organization and track perturbation-associated programs. (a) Workflow for breast-cancer single-cell benchmarks and Tahoe cell-line perturbation analysis. (b) Configured biological-program memberships or comparator gene sets/factors: scKNIFE benchmark v1, 7,521; in-house v2, 7,668; SPECTRA, 184; expiMap, 843; Slalom, 118. (c) Recorded training times for one 20,000-cell run per method and cohort in Bassez cohorts 1 and 2. (d–g) scKNIFE UMAPs for Bassez cohorts 1 and 2, Shiao and Wu TNBC; (h–k) corresponding UMAPs colored by author cell-type annotations. (l) ARI, NMI and AMI for shared-cell intersections in Bassez cohorts 1 and 2 and Wu TNBC. (m,n) MAGIC-imputed *PIK3R1* expression and PI3K–AKT–mTOR scores after afatinib in 60,743 Tahoe cells, shown for scKNIFE and Scanpy. (o,p) MAGIC-imputed *TP53* expression and apoptosis scores after doxorubicin in 64,064 Tahoe cells, shown for scKNIFE and Scanpy; Pearson r and Spearman *ρ* values are shown for each cell-level comparison.

We asked whether biologically informed latent representations retain the cellular organization of breast-cancer samples. For each method, we clustered cells using its learned cell-by-factor scores and compared the resulting partitions with the original study’s cell-type annotations on matched cells (Figure 2d-l; Supplementary Figure S2). In scKNIFE, clustering used the H-derived cell representation. The knowledge graph included gene sets representing major cell types; after clustering, we summarized cluster-level scKNIFE cell-identity and marker-node activities and used the highest-scoring available feature to characterize each cluster. Thus, the factorization did not use per-cell author labels as targets, although major-cell-type gene sets were part of the graph. The scKNIFE H representation grouped recognizable epithelial, immune, and stromal populations across Bassez cohorts 1 and 2 (also labeled A and B), Shiao, and Wu TNBC ^1–3^. For benchmark comparisons, cluster number was selected using the reference class count and silhouette scores, as described in Methods. On exact shared-cell intersections for the three datasets with all four methods, scKNIFE ARIs were 0.731, 0.794, and 0.678 in Bassez 1, Bassez 2, and Wu TNBC, respectively (Figure 2l). Corresponding SPECTRA ARIs were 0.738, 0.846, and 0.767; expiMap ARIs were 0.465, 0.883, and 0.636; and Slalom ARIs were 0.279, 0.425, and 0.424. The starred SPECTRA entries used cell-type supervision (Figure 2l). These comparisons describe alignment between factor-derived clusters and study annotations, rather than supervised per-cell label prediction. Together, the results show that scKNIFE retains recognizable cell-type organization while providing activity scores for downstream functional analyses.

We also compared recorded training times for the same 20,000-cell Bassez runs. scKNIFE took 1.44 and 2.13 h in cohorts 1 and 2, respectively, compared with 12.49 and 13.14 h for SPECTRA, 14.72 and 15.45 h for expiMap, and 7.32 and 11.43 h for Slalom (Figure 2c). Each value reflects one run per method and cohort under the retained implementation settings. Final scKNIFE parameters and tuning records are documented in Supplemental Tables 3A-C.

We then assessed how consistently independent cohort fits represented graph-node loadings (Supplementary Figure S1). Linear centered-kernel alignment of normalized *W* matrices was highest between Bassez 1 and 2 (0.895), with values of 0.669-0.826 for the remaining dataset pairs. Effective ranks were 22.21 of 40 factors in Bassez 1, 22.90 of 40 in Bassez 2, 56.80 of 80 in Shiao, and 15.78 of 30 in Wu. Pooled *W* -space embeddings by node type and unsupervised cluster provided a complementary view. Together, the alignment and rank estimates indicate shared loading-space structure alongside cohort-specific effective dimensionality.

Finally, we asked whether inferred program scores tracked expression of genes associated with the expected perturbation responses in the Tahoe compendium. MAGIC-imputed *TP53* expression correlated with scKNIFE apoptosis activity among 64,064 doxorubicin-exposed cells (Pearson *r* = 0.646; Spearman *ρ* = 0.537; Figure 2o-p); corresponding Scanpy correlations were *r* = 0.626 and *ρ* = 0.398. Among 60,743 afatinib-exposed cells, *PIK3R1* expression correlated with scKNIFE PI3K-AKT-mTOR activity (*r* = 0.765; *ρ* = 0.703), compared with *r* = 0.745 and *ρ* = 0.696 for Scanpy scoring (Figure 2m-n). These comparisons relate inferred scores for well-characterized apoptosis and PI3K-AKT-mTOR programs to expression of biologically relevant genes. A related expression-based interpretation was used in SPECTRA, where factor scores were compared with MAGIC-imputed *IFNGR1* expression in macrophages ^5^.

Together, these benchmarks show that scKNIFE represents broad biological knowledge, retains recognizable cellular organization across cohorts, and links inferred programs to relevant gene expression. We next examined whether this representation could resolve patient-level metabolic variation associated with clinical response, T-cell expansion, and treatment.

### scKNIFE resolves baseline tumor programs and treatment-associated functional changes across breast-cancer compartments

We next examined whether scKNIFE could identify functional programs associated with treatment outcomes across two independent breast-cancer studies. Shiao profiled serial TNBC biopsies during pembrolizumab-based treatment, distinguishing clinical responders (R) from non-responders (NR) ^3^. Bassez profiled breast tumors before and during anti-PD-1 treatment, distinguishing patients with subsequent T-cell clonotype expansion (E) from those without expansion (NE) ^2^. These complementary endpoints allowed us to examine baseline tumor features and treatment-associated changes across malignant, immune and stromal compartments.

We first compared scKNIFE with scCellFie, which infers metabolic-task activities through gene-protein-reaction rules ^8^. Pooled-cell treatment screens identified substantial overlap between the methods in Bassez cohort B and Shiao (Figure 3a-d). At a BH-adjusted *P <* 0.05 threshold, the methods shared 766 and 533 cell-type-task associations in the Bassez E and NE groups, respectively, and 863 and 984 in Shiao responders and non-responders. Shared associations represented more than half of the combined set in each group, with matching directions for 57-80% of shared associations. scKNIFE also identified additional associations across all four groups. This correspondence motivated closer examination of selected programs at the patient level.

**Figure 3.**
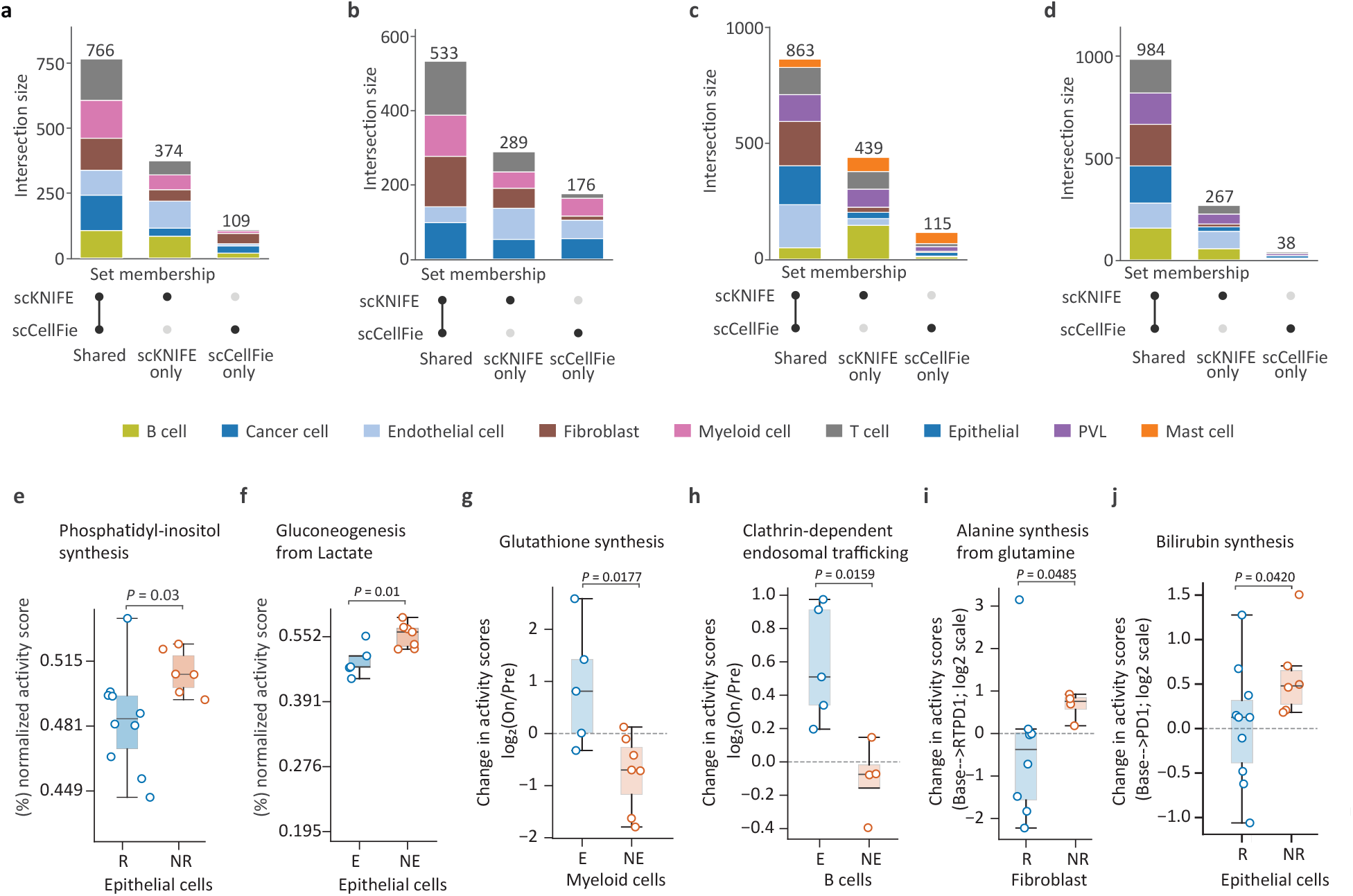
Baseline and treatment-associated metabolic activity in external breast-cancer cohorts. (a–d) UpSet plots of scKNIFE and scCellFie cell-type–task associations passing method-specific BH-adjusted *P <* 0.05 in Bassez cohort 2 clonotype-expanded (E) and non-expanded (NE) groups and Shiao responders (R) and non-responders (NR). (e) Baseline tumor/epithelial phosphatidylinositol-synthesis activity in Shiao R (n=10) and NR (n=6) patients. (f) Baseline tumor/epithelial gluconeogenesis-from-lactate activity in Bassez cohort 1 E (n=5) and NE (n=7) patients. Boxes show medians and interquartile ranges; points show patient medians. (g–j) Patient-level log_2_(treatment/baseline) activity changes: (g) Myeloid glutathione synthesis and (h) B-cell clathrin-dependent endosomal trafficking in Bassez cohort 1; (i) Fibroblast alanine synthesis from glutamine at RTPD1 and (j) tumor-cell bilirubin synthesis at PD1 in Shiao. Points show patients meeting the cell-count threshold at both timepoints; boxes show medians and interquartile ranges, whiskers extend to 1.5 interquartile ranges, and the dashed line marks zero change.

Pretreatment tumor programs highlighted lipid and lactate-associated metabolism as features linked to subsequent outcomes. In Shiao, inferred phosphatidylinositol-synthesis activity was higher in patients who later did not respond to treatment (nominal patient-level *P* = 0.0321; Figure 3e). Phosphatidylinositol composition has been associated with immune-checkpoint gene expression in breast cancer ^15^, providing biological context for this baseline lipid-associated difference. In Bassez, tumor gluconeogenesis-from-lactate activity was higher in patients without subsequent clonotype expansion (cohort A, nominal *P* = 0.0126; Figure 3f), with the same direction in the smaller cohort B. The established capacity of breast-cancer cells to use extracellular lactate supports the relevance of this inferred lactate-linked program ^16^. Patient concordance and cellular activity maps further characterize these baseline differences (Supplementary Figure S3A, panels A-B; Supplementary Figure S3B, panels A-C).

During treatment, clonotype expansion was accompanied by changes in myeloid redox metabolism. Glutathione-synthesis activity increased in E patients and decreased in NE patients in Bassez cohort A (nominal patient-level *P* = 0.0177; Figure 3g), with the same directional pattern in the cohort-B heatmap (Supplementary Figure S3A, panels E-F). Glutathione contributes to redox buffering, while experimental studies have linked redox regulation to macrophage PD-L1 expression and immunoregulatory behavior ^17^. These observations identify myeloid redox adaptation as a functional feature accompanying the different clonotype-expansion states.

B-cell trafficking provided a complementary immune-associated program. Clathrin-dependent endosomal trafficking increased in E patients but declined slightly in NE patients (nominal patient-level *P* = 0.0159; Figure 3h). Clathrin participates in B-cell receptor internalization ^18^, connecting this inferred activity to receptor handling. Patient-level GO analyses also identified greater increases in B-cell immunological-synapse and memory-T-cell-differentiation-related programs in E patients after BH correction (Supplementary Figure S3A, panel C; corresponding cohort-B comparisons in panel D). Together, these findings connect clonotype expansion with broader remodeling of B-cell trafficking and immune-interaction programs.

In Shiao, clinical response was associated with contrasting stromal nutrient-related trajectories. Fibroblast alanine-synthesis-from-glutamine activity decreased in responders but increased in non-responders during pembrolizumab plus radiation (nominal patient-level *P* = 0.0485; Figure 3i). Evidence for glutamine-mediated fibroblast-TNBC interactions ^19^ and stromal alanine support of pancreatic tumor metabolism ^20^ provides a biological rationale for examining this program. The opposing trajectories place fibroblast amino-acid handling among the stromal processes associated with treatment response.

Tumor bilirubin synthesis revealed a further connection to immune regulation. During pembrolizumab alone, the median patient-level increase was larger in non-responders than responders (nominal *P* = 0.0420; Figure 3j). Published breast-cancer experiments show that tumor-derived bilirubin can alter macrophage function and increase macrophage-mediated T-cell suppression ^21^. The stronger inferred program in non-responders therefore nominates heme-associated metabolism as a potential link between tumor metabolic state and the immune context of response.

Pooled-cell screens supported the within-group directions for glutathione synthesis, endosomal trafficking and alanine synthesis after BH adjustment, and the bilirubin increase in non-responders. These tests complement the patient-level comparisons of changes between groups (Supplemental Table 3D), while Supplementary Figure S3B, panels D-G, localizes the on-treatment activities. Together, these exploratory patient-level associations connect baseline tumor metabolism with treatment-associated functional changes across immune and stromal populations. We next examined these relationships within a paired TNBC atlas.

### A knowledge-guided TNBC representation retains cellular identity and patient variation

We assembled an atlas of 69,391 cells from 15 specimens collected from seven patients with TNBC (P1-P7). All seven contributed biopsy and surgery specimens; P3 also contributed a specimen from resistant relapse (Figure 4a). The specimens differed in treatment, procedure, metastatic status, and recorded disease site (Figure 4b; Supplemental Table 4A). The proportions of the 12 original annotated cell classes varied across samples (Supplementary Figure S4a), and the atlas included substantial epithelial, immune, and stromal compartments (Supplementary Figure S4b), reflecting the diversity of breast-cancer ecosystems described by Wu et al. ^1^.

**Figure 4.**
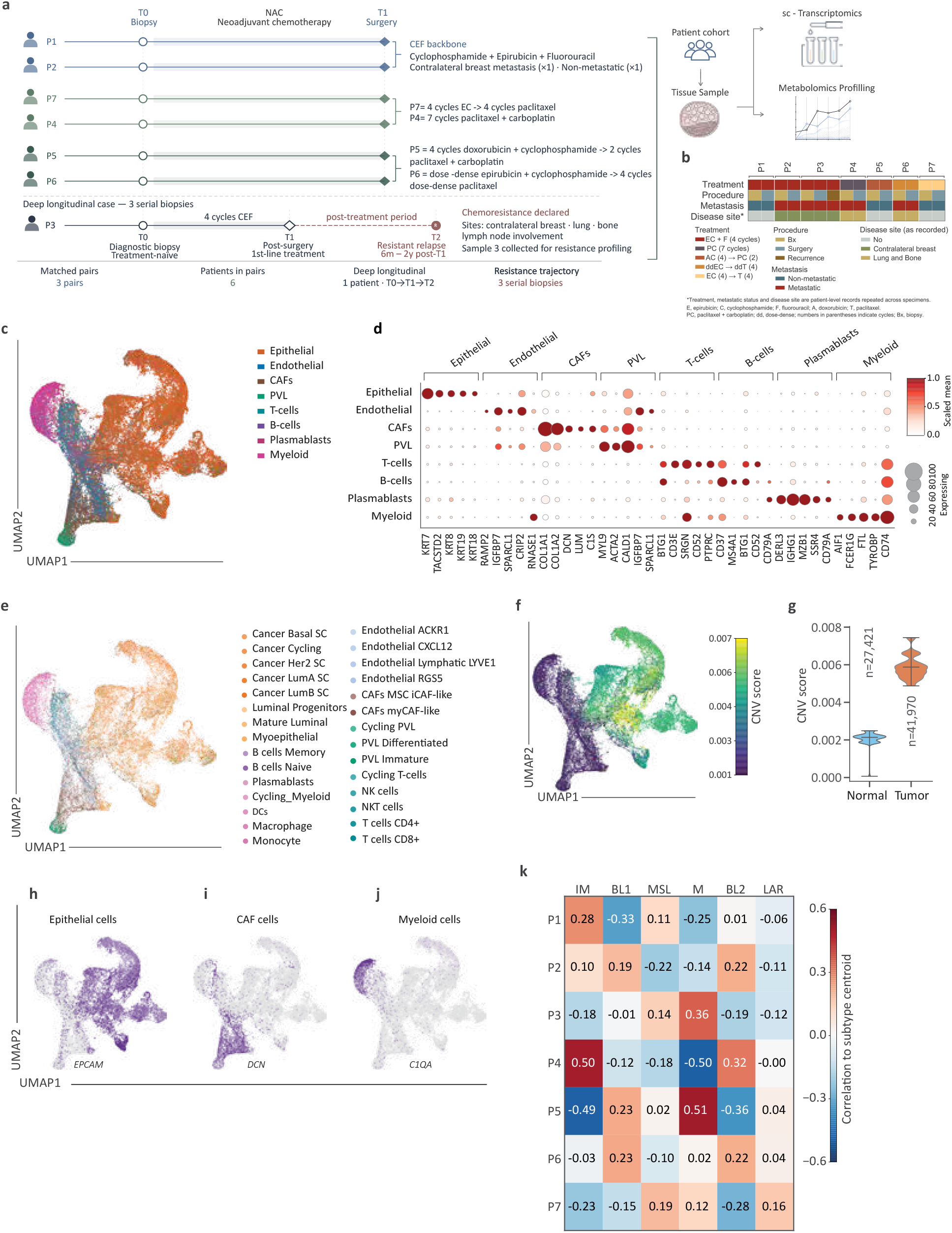
A seven-patient TNBC atlas retains broad and fine cellular identities in the H-derived representation. (a) Sampling and treatment schematic for seven pseudonymized patients (P1–P7). All seven patients contributed biopsy (T0) and surgery (T1) specimens; P3 also contributed a resistant-relapse specimen (T2). The schematic shows six patients in three two-patient display groups and P3 separately; all seven have biopsy–surgery pairs. (b) Patient-level clinical annotations repeated across specimens: treatment, procedure, metastatic-status and recorded disease-site annotations. The atlas contains 15 specimens. (c) SingleR major-label predictions against the Wu breast-cancer reference for all 69,391 cells on the saved H-derived UMAP. The eight broad display groups are Epithelial, Endothelial, CAFs, PVL, T-cells, B-cells, Plasmablasts and Myeloid; cancer and normal epithelial reference calls are combined in the Epithelial display label. (d) Five Wu-reference markers per broad group across the same eight groups (40 marker positions). Dot size indicates the percentage of cells with nonzero expression; white-to-red color indicates mean log1p-normalized expression scaled separately for each gene. (e) Cell-level full-Wu SingleR minor-label predictions on the same H-derived UMAP after default pruning. All 29 reference minor identities are keyed by color; 72 cells have ambiguous calls. Major and minor SingleR tiers were assigned independently. (f) Expression-derived CNV score on the H-derived UMAP; the display scale is clipped at the second and 98th percentiles. (g) Distributions of the same saved CNV scores in the provisional CNV-low/normal (*n* = 27, 421) and CNV-high/tumor (*n* = 41, 970) groups. CNV scores and status are transcriptome-derived annotations; the saved scores are assigned at CNV-cluster level and repeated across member cells. (h–j) *EPCAM, DCN* and *C1QA* expression, respectively, on the H-derived UMAP. (k) Heat map of patient-level correlation coefficients between epithelial profiles and six TNBCtype subtype centroids for P1–P7. Columns show the immunomodulatory (IM), basal-like 1 (BL1), mesenchymal stem-like (MSL), mesenchymal (M), basal-like 2 (BL2) and luminal androgen receptor (LAR) centroids; tiles display the correlation coefficients.

SingleR provided cell-level reference annotations for the 69,391-cell atlas using the full Wu breast-cancer reference. Major and minor identities were predicted independently for each cell and displayed on the saved H-derived UMAP (Figure 4c,e). The nine major reference labels were represented as eight broad groups by combining cancer and normal epithelial predictions into Epithelial: epithelial (*n* = 36, 963), cancer-associated fibroblasts (*n* = 9, 436), myeloid (*n* = 6, 241), T cells (*n* = 4, 921), perivascular-like cells (*n* = 3, 767), endothelial (*n* = 3, 493), plasmablasts (*n* = 3, 191), and B cells (*n* = 1, 379; Figure 4c; Supplemental Table 4B). The minor-label analysis retained 29 reference identities after pruning, with 72 cells classified as ambiguous (Figure 4e). Reference-marker expression across the broad groups is shown in Figure 4d and Supplemental Table 4D. The original 12-class annotation provided a complementary view through its 59-gene marker plot and atlas UMAP (Supplementary Figure S4c,d; Supplemental Table 4E). Together, the reference labels and marker expression show that the H-derived representation retained the cohort’s major epithelial, immune, and stromal organization.

Supplementary Figure S4g displays two reference-derived marker positions for each minor identity across all 29 groups (Supplemental Table 4E). These expression patterns place the fine-grained predictions in biological context, including *KRT14* in myoepithelial cells, *COL1A1* and *COL3A1* in myCAF-like fibroblasts, *CD3E* and *CD8A* in CD8-positive T cells, and *LYZ* and *VCAN* in monocytes. Cell-level overlap between the independently predicted major and minor SingleR labels is tabulated in Supplemental Table 4C.

The adjacent CNV-score map added tumor-associated structure to this cell-identity view (Figure 4f). The provisional CNV-high/tumor group contained 41,970 cells, whereas the CNV-low/normal group contained 27,421, with their score distributions separated in Figure 4g (Supplemental Table 4F). Supplementary Figure S4e,f shows the corresponding 26 expression-derived CNV clusters and provisional status on cell-level embeddings. *EPCAM, DCN* and *C1QA* localized to epithelial-rich, fibroblast-rich and myeloid-rich atlas regions, respectively (Figure 4h-j). Read together, the reference labels, marker expression and transcriptome-derived CNV patterns resolve cellular compartments and tumor-associated variation on the same H map, in line with single-cell breast-cancer profiling approaches ^1,22^.

To place this cellular heterogeneity within established TNBC transcriptional classes, we examined patient-level correlations with the six TNBCtype subtype centroids (Figure 4k). P1 and P4 showed their strongest correlations with the immunomodulatory signature (*r* = 0.28 and 0.50), whereas P3 and P5 correlated most strongly with the mesenchymal signature (*r* = 0.36 and 0.51). P7 showed a weaker preference for the mesenchymal stem-like signature (*r* = 0.19). Immune and stromal contributions to the immunomodulatory and mesenchymal stem-like signatures have been established previously ^23^, placing these patient-level profiles in the context of the cellular diversity resolved by the atlas. We next examined epithelial cells separately to identify functional programs underlying variation among patients and between biopsy and surgery (Figure 5).

**Figure 5.**
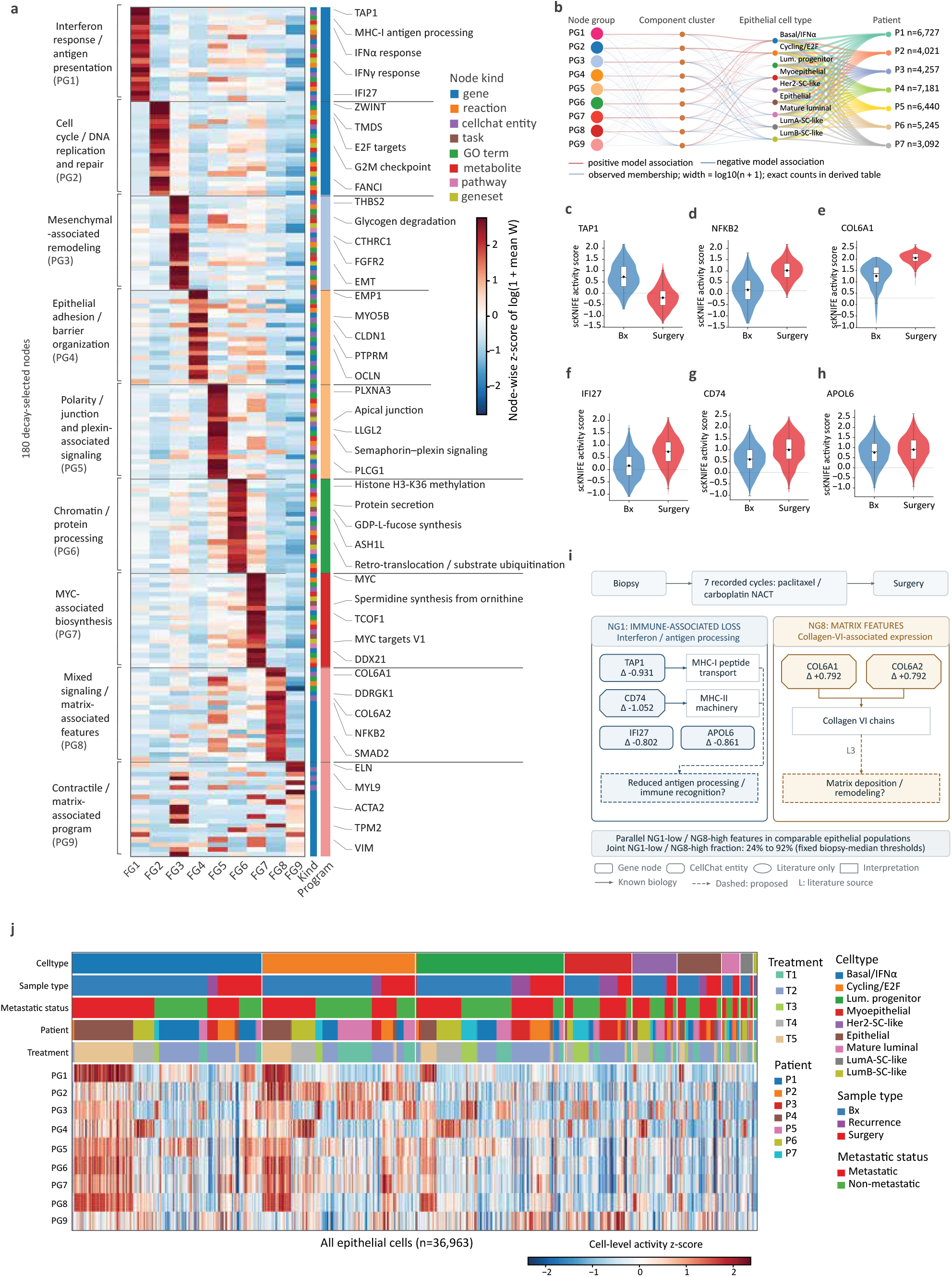
scKNIFE resolves epithelial programs and a coordinated residual state in P4. (a) Node-wise z scores of log(1 + mean W) for 180 representative nodes from nine W-derived groups (PG1–PG9). Twenty globally unique nodes per group were selected by exponential decay (*τ* =5, cumulative mass=0.95); five per group are labelled. Cell-type and cell-state graph nodes were excluded before downstream W clustering and node selection. (b) Network connecting node groups to component clusters and epithelial/state groups through signed model associations. The final layer shows 63 observed epithelial/state-group-to-patient count links totaling 36,963 cells; line widths represent log_10_(n+1). (c–e) Inferred *TAP1* (c), *NFKB2* (d), and *COL6A1* (e) node activity in P4, comparing biopsy and surgery after matching by epithelial cell type and RNA depth (434 cells per specimen). P4 had seven recorded cycles of paclitaxel/carboplatin between specimens and subsequently recorded lung and bone metastases. (f–h) Corresponding *IFI27* (f), *CD74* (g), and *APOL6* (h) distributions in P1 (1,681 matched cells per specimen), who had no recorded metastasis. Violin boxes show medians and quartiles. Node activity is reconstructed from the aligned H and W matrices, log1p-transformed and standardized across all 36,963 displayed epithelial cells; the cells at the two timepoints are distinct sampled cells. (i) Schematic of P4’s paired-sample pattern. Mean matched-cell PG1 activity decreased by 0.809 global z-score units and PG8 increased by 1.320 (+0.15*→*+1.47); the schematic labels these groups NG1 and NG8. Across all P4 epithelial cells, mean PG8 instead fell from +1.41 to +1.03 (Supplementary Figure S5J); the fraction classified PG1-low/PG8-high rose from 24% (104/434) at biopsy to 92% (400/434) at surgery using fixed biopsy-median thresholds. Solid arrows indicate established molecular functions; dashed boxes indicate proposed biological consequences. (j) Reconstructed PG1–PG9 activity across all 36,963 epithelial cells, grouped by nine provisional epithelial/state annotations spanning 21 inherited Leiden 0.8 clusters. Annotation tracks show cell type, sample type, metastatic status, patient, and treatment. Panels c–i are descriptive within-patient comparisons; Supplementary Figure S5J,K shows the all-cell and matched-cell estimator sensitivity.

### scKNIFE resolves epithelial programs and a coordinated residual state in P4

Within the paired TNBC atlas, we asked whether aligned scKNIFE factors could connect shared epithelial identities to functional states that vary among patients and between biopsy and surgery. The analysis displayed 36,963 cells assigned to the broad epithelial group by Wu-reference SingleR annotation, using aligned H and W matrices from the 50-factor epithelial fit. That fit used 43,574 cells selected under the earlier annotation for the TNBC study cohort. The 50 factors were organized into nine W-derived program groups (PG1-PG9; Figure 5a; Supplemental Tables 5A,B), corresponding to NG1-NG9 in retained analysis files. The schematic in Figure 5i retains NG labels. The heatmap of 180 representative nodes resolved interferon and antigen-presentation features (PG1), cell-cycle and DNA-replication features (PG2), a *MYC*-associated biosynthesis program containing spermidine synthesis from ornithine (PG7), mixed signaling and matrix-associated features (PG8), and a contractile program containing *ACTA2* and *MYL9* (PG9). These themes parallel the immune-rich and matrix-rich breast-cancer phenotypes reported by Asleh et al. ^24^, cycling heterogeneity in TNBC described by Karaayvaz et al. ^25^, and the relationship between *MYC* and polyamine metabolism investigated in TNBC ^26^. The inferred W neighborhoods organize these features into programs that can be followed from their constituent nodes to epithelial states and individual patients.

The program-group-to-component-cluster-to-epithelial-state network linked PG1 most strongly to basal/IFN*α* cells, PG2 and PG7 to cycling/E2F cells, and PG9 to myoepithelial cells (Figure 5b). The first two edge layers represent signed model associations; the 63 patient-facing links represent observed cell counts and sum to all 36,963 epithelial cells (Supplemental Table 5F). This alignment with recognizable breast epithelial states is consistent with the heterogeneity mapped by Wu et al. ^1^. Patient coverage and cell-weighted versus equal-patient summaries in Supplementary Figure S5A-C show the distribution underlying these relationships. The PG7 metabolic-task plots and patient summaries provide additional program context (Supplementary Figure S5D-I; Supplemental Table 5G).

We then examined cell-type- and RNA-depth-matched biopsy and surgical epithelial populations from two patients. Reconstructed node activities were log1p-transformed and standardized across the 36,963 displayed epithelial cells; negative scores therefore indicate activity below the atlas mean. In P4 (434 cells per specimen), median inferred *TAP1* activity fell from 0.698 to *−*0.209 (mean change *−*0.931; Figure 5c), whereas *NFKB2* rose from 0.053 to 0.981 (+0.939; Figure 5d) and *COL6A1* from 1.079 to 1.799 (+0.792; Figure 5e). *TAP1* contributes to MHC-I antigen processing, and Rädler et al. observed lower MHC-I and interferon expression in paired basal-like residual TNBC specimens after neoadjuvant chemotherapy ^27^. The accompanying *NFKB2* and *COL6A1* changes nominate signaling and matrix-associated features in P4’s residual epithelial population: chemotherapy-exposed breast-cancer models have shown increased noncanonical NF-*κ*B components ^28^, and collagen VI can support breast-cancer-cell invasion experimentally ^29^. The three trajectories thus identify related features within a matched patient comparison (Supplemental Table 5C).

P1 (1,681 matched cells per specimen) showed another treatment-stage pattern (Figure 5f-h): median *IFI27* activity rose from 0.141 to 0.737, *CD74* from 0.595 to 1.021, and *APOL6* from 0.798 to 0.916. Their concordant directions suggest an interferon- and antigen-presentation-associated epithelial state. Interferon-stimulated tumor-cell states have been observed in TNBC, including residual disease after chemotherapy ^30^; Forero et al. demonstrated *CD74* and MHC-II pathway expression in TNBC tumor cells ^31^; and *APOL6* responds to interferon in breast-cancer-cell experiments ^32^. These paired examples reveal distinct patient-specific combinations of immune-associated and signaling/matrix-associated features (Supplemental Table 5C).

Figure 5i summarizes these P4 node-level changes. In matched epithelial populations, mean PG1 activity fell by 0.809 global z-score units, while mean PG8 activity rose by 1.320 (from +0.15 to +1.47). This differs from the all-epithelial estimate, where PG8 fell from +1.41 to +1.03 between biopsy and surgery (Supplementary Figure S5J); the residual-state pattern therefore applies to the matched populations. Using fixed biopsy-median thresholds, the fraction classified PG1-low/PG8-high rose from 24% (104/434 cells) at biopsy to 92% (400/434) at surgery. *CD74, IFI27*, and *APOL6* also decreased with PG1 (*−*1.052, *−*0.802, and *−*0.861; Supplemental Table 5D), while *COL6A2, SMAD2*, and *DDRGK1* increased with PG8 (+0.792, +1.262, and +0.767). Prior studies link paclitaxel exposure to increased TGF-*β* signaling in TNBC models ^33^ and *DDRGK1* to endoplasmic-reticulum homeostasis ^34^. These findings nominate immune-associated and tissue-interaction features of P4’s residual epithelial population. P4’s subsequent lung and bone metastases provide clinical context but do not establish a link between this state and metastasis.

Across all 36,963 epithelial cells, the reconstructed activity heatmap placed PG2 and PG7 in cycling/E2F cells (8,281 cells) and PG9 in myoepithelial cells (3,653 cells; Figure 5j; Supplemental Table 5F). The equal-patient summaries retained these program-state relationships (Supplementary Figure S5C; Supplemental Table 5E). Thus, the atlas places the P4 residual-state pattern and the P1 interferon-associated pattern within a broader landscape of epithelial identity and patient variation.

The matched and all-cell summaries describe different sampled populations (Supplementary Figure S5J,K). In P1, mean all-cell PG1 activity rose from +0.14 to +0.91, consistent with the interferon-associated node changes (Supplementary Figure S5K). The P4 matched-population shift and P1 interferon-associated pattern illustrate distinct patient-specific epithelial changes. We then examined metabolic activity across compartments alongside paired tissue measurements.

### Cell-resolved metabolic activities connect tissue pathway enrichment to myeloid lipid remodeling

To connect cellular predictions with measured tissue context, we integrated paired LC-MS profiles and scKNIFE activities from 14 biopsy and surgery specimens from seven patients. Of 21 treatment-associated tissue pathway candidates, 18 source terms mapped to 14 distinct Human-GEM pathways represented in the cell-based scores; five candidates passed Holm-adjusted Gamma testing: selenoamino-acid, purine, methionine/cysteine, aspartate/asparagine, and glycolysis/gluconeogenesis metabolism. This mapping enabled follow-up of tissue-level signals in specific cellular compartments (Figure 6a,b; Supplemental Table 6A). No pathway passed Holm correction in the separate metastatic-status analysis (Supplementary Figure S6A,B). These enrichments provide pathway context for the cell-resolved hypotheses, not independent validation of scKNIFE activities.

**Figure 6.**
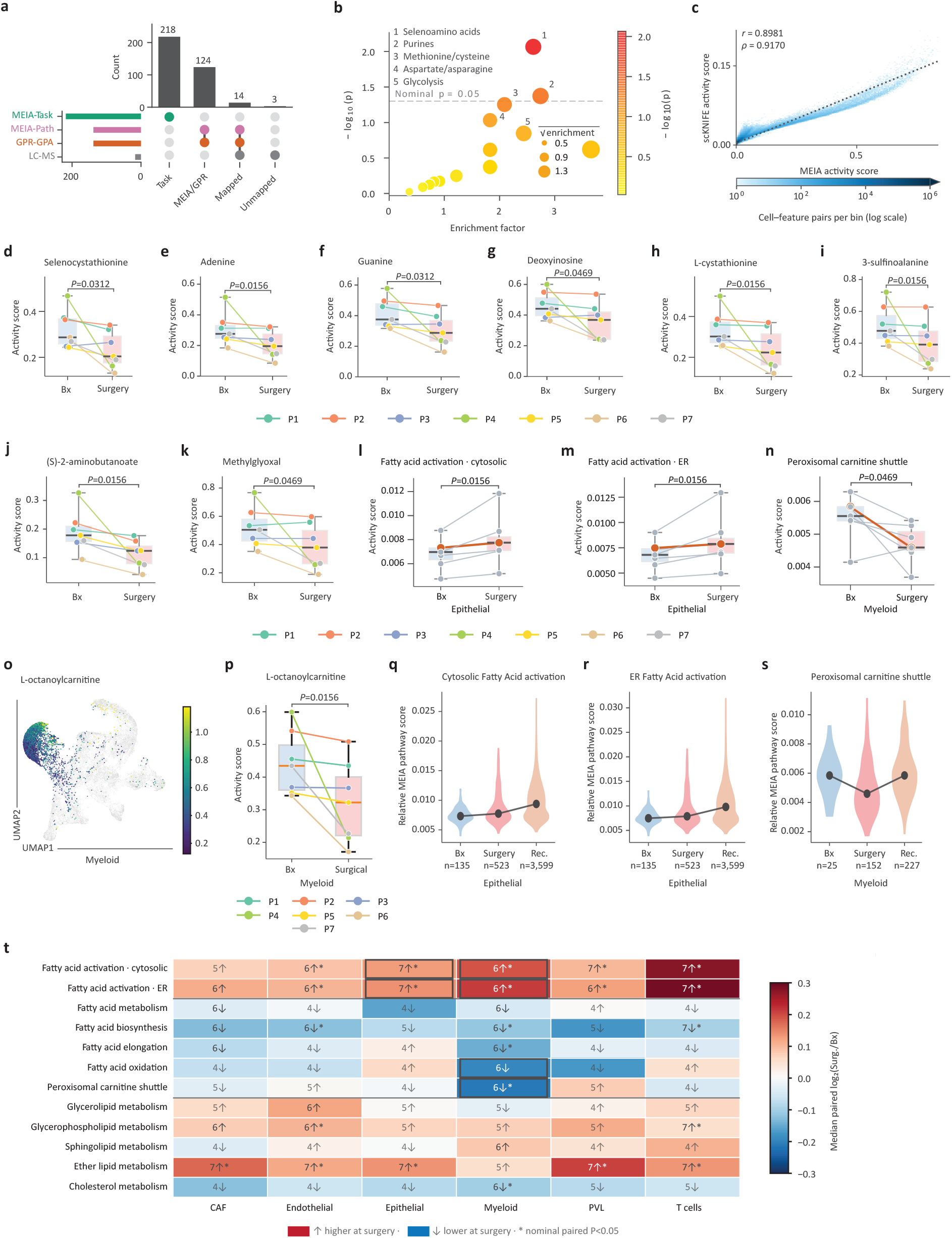
Tissue metabolic pathway context and patient-resolved inferred activity after neoadjuvant treatment in TNBC. (a) Coverage of 21 LC–MS candidate pathways by MEIA and GPR-GPA scores. (b) Surgery-versusbiopsy LC–MS pathway enrichment; each point represents a candidate pathway and the vertical axis shows *−* log_10_(Fisher *P*). (c) MEIA and direct scKNIFE task-score concordance across 69,390 shared cells and 218 tasks (15,127,020 cell–task pairs); each task block is normalized within each cell, and Pearson *r* = 0.8981 and Spearman *ρ* = 0.9170 are shown. (d–k) Paired biopsy and surgery Myeloid activity scores for selenocystathionine (d), adenine (e), guanine (f), deoxyinosine (g), L-cystathionine (h), 3-sulfinoalanine (i), (S)-2-aminobutanoate (j) and methylglyoxal (k) in seven patients. Boxes show patient medians and interquartile ranges, whiskers show the observed range, and points and lines identify paired patients; displayed *P* values are nominal exact two-sided paired Wilcoxon signed-rank tests. (l–n) Paired patient-level MEIA scores for Epithelial cytosolic fatty-acid activation (l), Epithelial endoplasmic-reticular fatty-acid activation (m) and Myeloid peroxisomal carnitine shuttle (n) in seven patients; boxes show medians and interquartile ranges, and lines connect paired values. Displayed *P* values are from exact two-sided paired Wilcoxon signed-rank tests. (o) Saved H-derived UMAP with 6,014 biopsy and surgery Myeloid cells colored by inferred L-octanoylcarnitine activity; grey points show all atlas cells, and the color scale spans the second to 98th percentiles. (p) Paired patient-level Myeloid L-octanoylcarnitine scores; points show patient medians and lines connect pairs, with exact two-sided paired Wilcoxon signed-rank *P* values. (q–s) P3 cell-level distributions at biopsy, surgery and recurrence for Epithelial cytosolic fatty-acid activation (q), Epithelial endoplasmic-reticular fatty-acid activation (r) and Myeloid peroxisomal carnitine shuttle (s); dots mark specimen medians, lines connect the three medians, and labels show cell counts. (t) Median patient-level log_2_(surgery/biopsy) ratios for 12 inferred Human-GEM lipid pathways across six cell compartments. Tiles show ratio direction and patient counts; bold symbols mark six or seven concordant pairs. Asterisks mark nominal *P <* 0.05 from exact two-sided paired Wilcoxon signed-rank tests on raw patient-median surgery-minus-biopsy score differences; Benjamini–Hochberg *q* values use the full 828-test pathway-by-cell family. For panels d–n and p, points represent specimen-level patient medians and lines connect paired biopsy and surgery values from the same patient (seven pairs). Boxes span the 25th–75th percentiles, the central line marks the median, and whiskers span the observed range. Displayed *P* values are nominal, two-sided exact paired Wilcoxon signed-rank tests. In panel t, tile colors show median paired log_2_(surgery/biopsy) ratios, whereas asterisks mark nominal paired Wilcoxon tests on raw patient-median surgery-minus-biopsy score differences. Screen-wide adjusted values are reported in Supplemental Tables 6C–G. Panels q–s show distinct sampled cells from the single patient P3 and are not recurrence-cohort comparisons.

Metabolic evidence integration analysis (MEIA) and direct reconstructed scKNIFE task scores were strongly concordant across shared cells (Figure 6c; Supplemental Table 6B). Since both constructions use related transcriptomic and graph inputs, this agreement compares scoring approaches rather than providing independent validation. We assigned cells to six Wu-reference compartments by exact cell ID and summarized paired biopsy-surgery changes at the patient level (Supplementary Figure S6C-E).

Across the seven paired patients, LC-MS-linked myeloid scores for sulfur- and purine-associated nodes were generally lower at surgery (Figure 6d-k; Supplemental Tables 6C,H). The displayed nodes included selenocystathionine, adenine, guanine, deoxyinosine, L-cystathionine, two nodes linked through Human-GEM reactions to the aspartate/asparagine term, and methylglyoxal, linked to glycolysis/gluconeogenesis. Each score was lower in six or seven of the seven patient pairs. The directions were retained in a sensitivity analysis excluding P7, whose surgery specimen contained only nine myeloid cells (Supplemental Table 6C). These candidates arose from a 1,344-test screen, and none passed global Benjamini-Hochberg correction; they therefore represent exploratory, pathway-linked patterns.

An LC-MS-independent pathway screen showed contrasting lipid-related directions across compartments (Figure 6l-n; Supplemental Table 6D). Cytosolic and endoplasmic-reticular fatty-acid activation scores increased in all seven patients in epithelial cells, whereas the myeloid peroxisomal carnitine-shuttle score decreased in six of seven. None of the 828 pathway-cell comparisons passed Benjamini-Hochberg correction. Thus, the selected pathway scores suggest compartment-specific differences in lipid handling, rather than a uniform change across cell populations.

The myeloid L-octanoylcarnitine score connected this pathway-level pattern to a constituent metabolite node: inferred activity was lower at surgery in all seven patients (Figure 6o,p; Supplemental Tables 6E,F). L-carnitine and octanoyl-CoA scores showed a similar, though less consistent, direction (Supplemental Table 6E). The focused carnitine-node screen yielded no finding after multiple-testing correction. These scores describe the broader myeloid compartment and do not establish a macrophage-specific change.

P3 contributed the only recurrence specimen. In this patient, epithelial cytosolic and ER fatty-acid activation scores rose across biopsy, surgery, and recurrence, while the myeloid peroxisomal carnitine-shuttle score fell at surgery and returned near its biopsy level at recurrence (Figure 6q-s). This is a single-patient trajectory, not a recurrence-cohort comparison; the pathway-level return does not imply a corresponding rebound in the L-octanoylcarnitine node. In the broader lipid summary, myeloid cytosolic and ER fatty-acid activation increased in six of seven patients, while fatty-acid oxidation and the peroxisomal carnitine shuttle decreased in six of seven. Epithelial fatty-acid activation and ether-lipid metabolism increased in all seven pairs (Figure 6t; Supplemental Table 6G). Several pathway-compartment comparisons were nominally supported, but the screen-wide pattern remained exploratory.

Together, the paired analyses connect tissue-level pathway enrichment to candidate myeloid purine and sulfur-associated activities, then place broader lipid-related changes within specific cellular compartments and patient trajectories. The inferred scores prioritize hypotheses but do not measure metabolite abundance or flux. We next examined candidate communication between atlas compartments.

### Patient-resolved PDGF communication

We compared scKNIFE and CellChat across 69,319 joined cells assigned to the same eight broad SingleR groups (Figure 7a-d). scKNIFE combines inferred ligand and receptor activities with cell-type specificity and curated interaction weights; CellChat provides an expression-based comparison ^9^. We used the shared-cell set to assess method overlap before following a selected interaction into the 29 independently predicted, pruned minor SingleR identities and paired patient samples.

**Figure 7.**
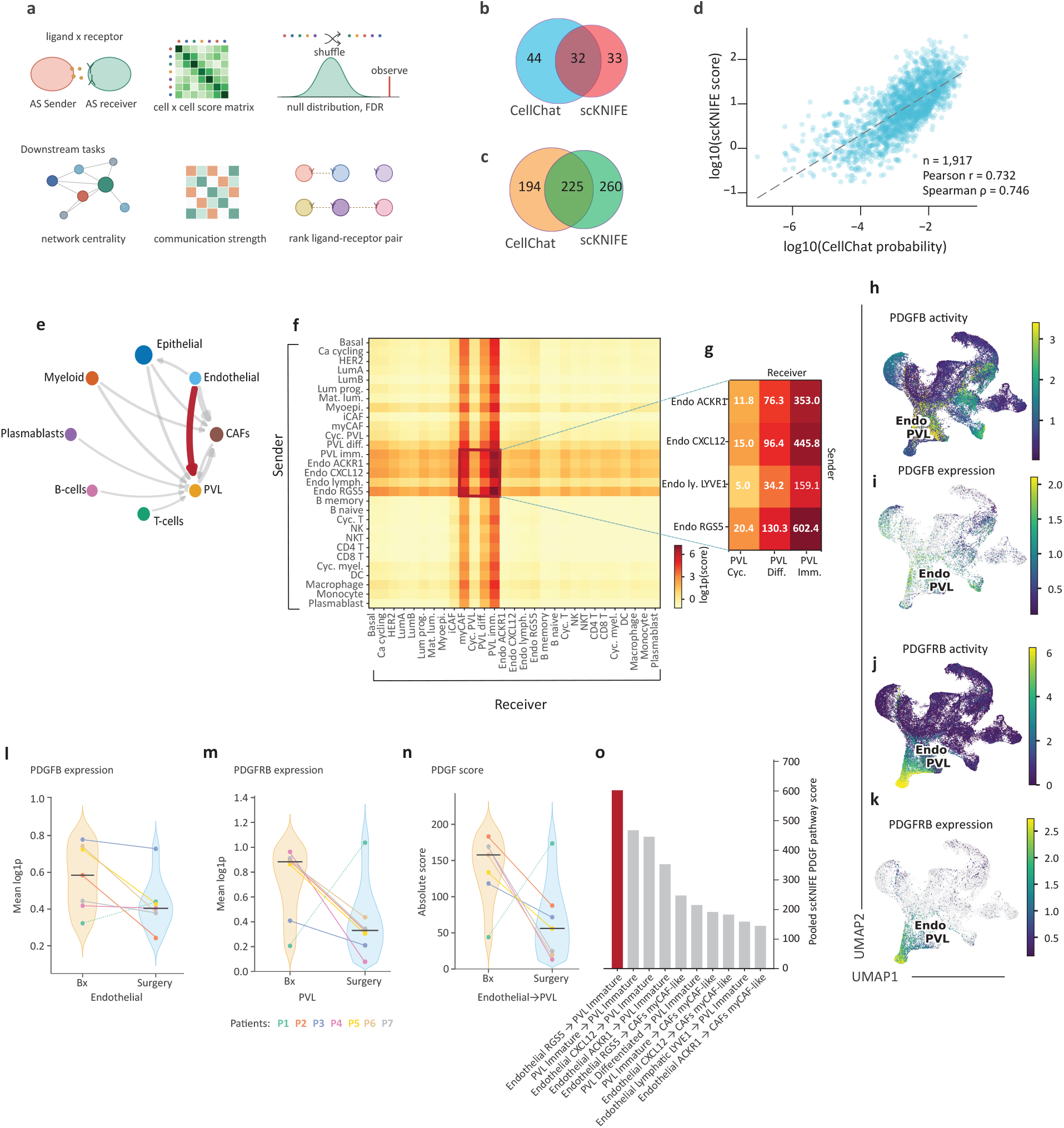
Cross-method concordance and patient-resolved PDGF communication in TNBC. (a) Overview of the cell–cell communication analysis. (b) Filtered pathway overlap between CellChat and scKNIFE. (c) Overlap of harmonized ligand–receptor keys. (d) Log-scale scores for 1,917 shared positive source–target/ligand–receptor records, with Pearson r and Spearman *ρ*. (e) Pooled eight-compartment PDGF network. Node colors identify compartments; arrowheads indicate sender-to-receiver direction, arrow width is scaled to the pooled scKNIFE PDGF score, and red highlights the selected Endothelial-to-PVL connection while gray indicates the other displayed connections. (f) Pooled 29*×*29 minor-cell sender–receiver PDGF matrix colored by log(1 + score). (g) Endothelial-sender/PVL-receiver slice of the matrix. (h–k) Inferred *PDGFB* activity, *PDGFB* expression, *PDGFRB* activity and *PDGFRB* expression on the atlas UMAP, each with an independent color scale. (l,m) Patient-level endothelial *PDGFB* and PVL *PDGFRB* expression at biopsy and surgery. (n) Paired patient-level major Endothelial-to-PVL PDGF scores; points and lines show matched patients. (o) Ten highest-scoring pooled minor-cell sender-to-receiver PDGF connections.

Under the saved method-specific filters, the methods shared 32 pathway labels (44 CellChat-only and 33 scKNIFE-only; Figure 7b; Supplemental Table 7A) and 225 harmonized ligand-receptor keys (194 CellChat-only and 260 scKNIFE-only; Figure 7c; Supplemental Table 7B). Among 1,917 positive ligand-receptor/source/target records present in both methods, log_10_ scores had Pearson *r* = 0.732 and Spearman *ρ* = 0.746 (Figure 7d; Supplemental Table 7C). The methods therefore shared some network features while retaining method-specific results.

We focused on PDGF because the network highlighted an Endothelial-to-perivascular-like (PVL) connection. In the eight-group network, Endothelial*→*PVL was a prominent PDGF connection (pooled score 109.34; Figure 7e; Supplemental Table 7D). The 29-by-29 minor-identity matrix resolved this broad pairing (Figure 7f), with Endothelial *RGS5→*PVL Immature as the highest-scoring edge in the Endothelial-sender/PVL-receiver block (Figure 7g; Supplemental Table 7D). *PDGFB*-*PDGFRB* contributed 83.5% of the selected major connection and 88.9% of the selected minor connection (Supplemental Table 7E). Figure 7h-k maps inferred *PDGFB* and *PDGFRB* activities alongside their measured log1p expression on the Figure 4 UMAP. This candidate pair is consistent with established endothelial PDGF-B recruitment of PDGFR*β*-positive pericyte precursors ^35^ and the contribution of PDGFR*β*-positive perivascular progenitors to tumor-vessel support ^36^.

The selected axis also varied across seven patients with paired biopsy and surgery specimens. Mean endothelial *PDGFB* expression was lower at surgery in six patients and higher in P1 (Figure 7l; Supplemental Table 7F). Mean PVL *PDGFRB* expression followed the same six-to-one patient direction (Figure 7m; Supplemental Table 7F); both expression panels are descriptive and have no separate P values. The absolute inferred major Endothelial*→*PVL PDGF score was also lower in six patients and higher in P1, with median values of 157.72 at biopsy and 56.04 at surgery (Figure 7n; Supplemental Table 7F). Panels l-n show the seven patient values at each stage, with points and connecting lines preserving the pairs. For this selected connection, the exact two-sided sign-flip test of mean patient log_2_(surgery/biopsy) gave nominal *P* = 0.109375 in the 2,080-test major screen (Supplemental Table 7G). The relative PDGF share is in Supplemental Table 7F but is not plotted. These data describe a patient-level treatment-stage pattern along a recognized endothelial-perivascular axis; the comparison did not pass multiple-testing correction. Eligibility required at least ten sender and receiver cells at both timepoints; P3 recurrence is excluded.

Endothelial *RGS5→*PVL Immature ranked first among the ten highest-scoring pooled minor-cell PDGF connections, with a score of 602.39 in the full 29 *×* 29 matrix (Figure 7o; Supplemental Table 7D). This longitudinal example was selected after the major pathway/source/target screen, in which none of 2,080 comparisons passed BH *q <* 0.05 (Supplemental Table 7G). The patient-level directions are therefore exploratory. Figure 7 follows this candidate from broad cell groups to minor identities and paired patient samples.

## Discussion

scKNIFE connects cellular organization to multiple levels of biological function through an expression-anchored graph factorization. Spectra and expiMap demonstrate the value of incorporating prior knowledge into interpretable gene-program models ^5,6^. scKNIFE represents genes and connected functional entities—including reactions, metabolites and signaling components—as nodes with loadings on the same latent factors. This organization allows a factor to be examined through its molecular composition, its distribution across cells and its variation among patients. Across the evaluated cohorts, the resulting representations retained recognizable cellular organization and supported analyses of epithelial programs, metabolic activities and candidate communication. The shared representation provides a common basis for following these relationships across biological scales.

The external-cohort analyses connected treatment-associated variation with distinct aspects of tumor and microenvironmental biology. Pretreatment tumor programs highlighted lipid- and lactate-associated metabolism, while changes during treatment involved myeloid redox regulation, B-cell trafficking, fibroblast amino-acid metabolism and tumor heme-associated activity. The biological interpretation therefore extended across the malignant, immune and stromal compartments. Myeloid glutathione synthesis and B-cell endosomal trafficking accompanied differences in T-cell clonotype expansion, whereas fibroblast alanine synthesis and tumor bilirubin synthesis varied with clinical response. Considered together, these findings place metabolic and immune functions within the cellular contexts relevant to each treatment endpoint.

The paired TNBC atlas extended this analysis to the diversity of cell states found within individual tumors ^1^. Epithelial programs in P1 and P4 followed distinct patterns between biopsy and surgery, showing how patient-specific organization contributes to the functional landscape of the cohort. P4’s PG8 pattern also varied with the epithelial populations included in the comparison. Examining the program across the full compartment and within comparable populations revealed how cellular composition and within-population activity contribute to the aggregate signal. This distinction gives biological meaning to patient heterogeneity by identifying both the populations carrying a program and the changes occurring within them.

The metabolic findings illustrate how scKNIFE brings tissue-level observations into a cellular framework. LC–MS pathway enrichment established a metabolic context, and related purine- and sulfur-associated activities were identified within the Myeloid compartment. The biopsy-enriched purine pattern is consistent with experimental evidence linking purine metabolism to pro-tumor macrophage states ^37^, providing a biological context for the broader myeloid observation. The lipid analysis resolved further differences among connected processes. Fatty acid activation scores increased in Myeloid and Epithelial cells, whereas myeloid oxidation and the peroxisomal carnitine-shuttle scores were higher at biopsy in six of seven patients. These distinct directions show why individual components of lipid metabolism carry more information than a single aggregate pathway label.

The carnitine-associated findings connect a pathway-level pattern to a related metabolite node. Inferred L-octanoylcarnitine activity was higher at biopsy in all seven patients, alongside predominantly biopsy-enriched peroxisomal carnitine-shuttle activity. CROT catalyzes reversible octanoyl-group transfer between CoA and carnitine ^38^, providing biochemical context for this relationship. Together, the inferred pathway and node activities identify carnitine-associated metabolism as a candidate component of the differences between biopsy and surgical myeloid populations.

The communication analysis extended this interpretation to interactions between cellular compartments. Comparison with CellChat contextualized the inferred communication patterns ^9^, while the endothelial–perivascular PDGF connection could be resolved into its dominant *PDGFB*–*PDGFRB* contribution, finer cellular identities and paired patient profiles. This molecular organization accords with the established role of endothelial PDGF-B in pericyte recruitment ^35^. Lower endothelial *PDGFB* expression, perivascular *PDGFRB* expression and inferred PDGF connection scores at surgery in six patients together suggest a treatment-stage difference in the transcriptional support for this vascular interaction. The biological meaning emerges from the correspondence between the signaling components, their cellular locations and their patient-level directions.

These analyses support a compartment-resolved account of treatment-associated tumor variation. The cellular metabolic and communication associations are exploratory observations from seven paired patients, with recurrence represented by P3, while the tissue pathway findings arise from a separate measurement and statistical analysis. The inferred scores prioritize hypotheses but do not measure metabolite abundance or flux. Within this scope, scKNIFE brings molecular relationships into the interpretation of cellular heterogeneity: tumor programs retain their patient context, metabolic patterns acquire a cellular location, and communication signals resolve into defined ligand–receptor relationships. This integration makes the biological organization underlying single-cell variation more explicit.

## Methods

### Key resources table

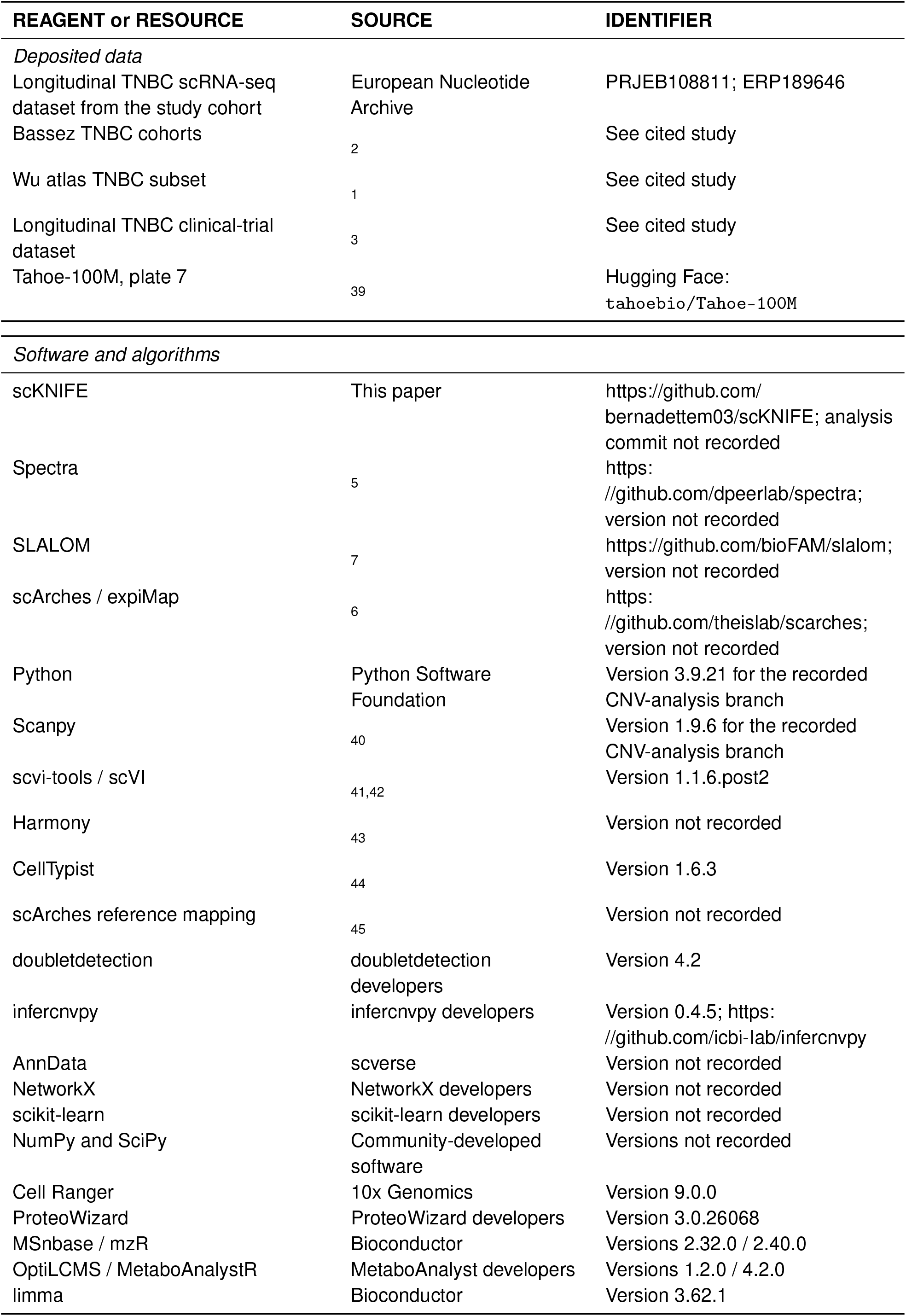
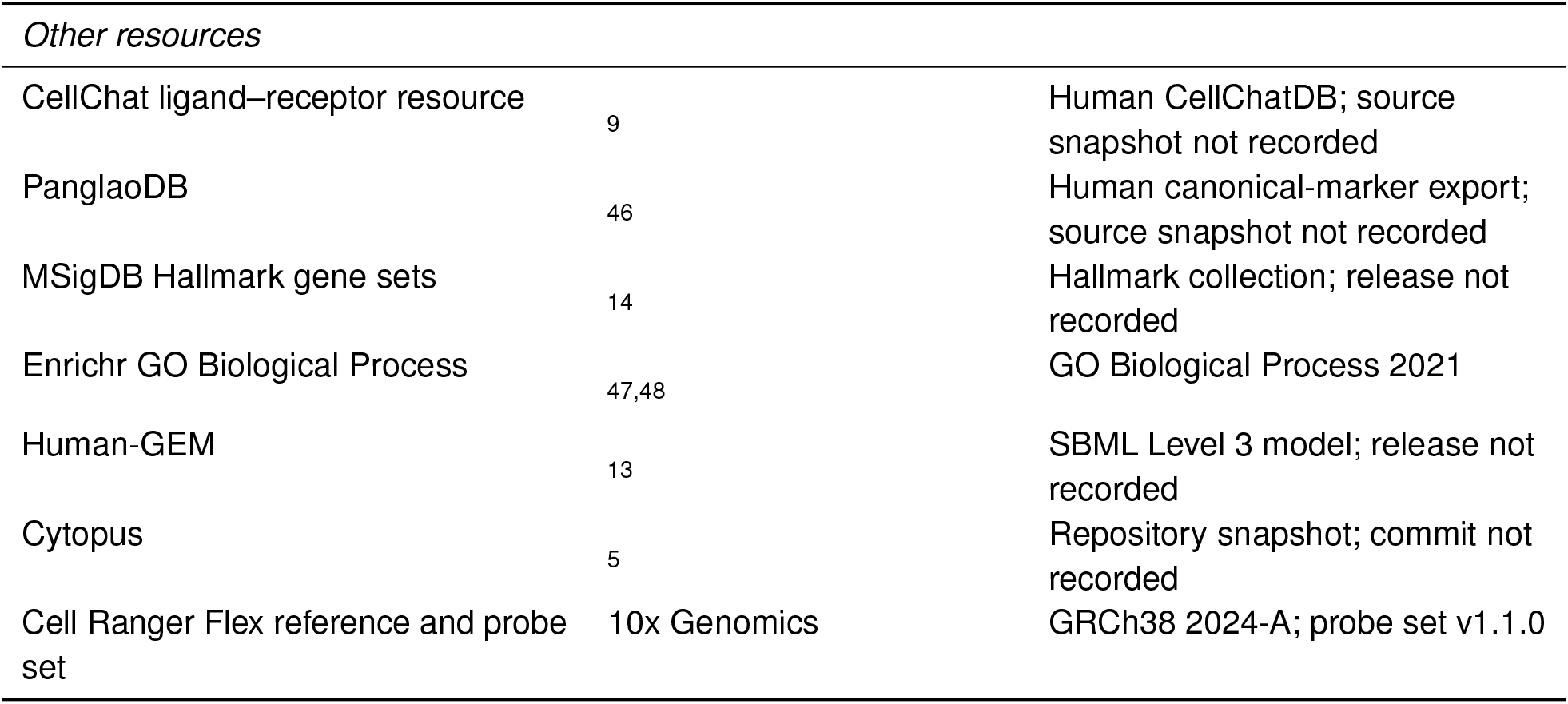

### Experimental model and study participant details

#### Study design, cohort, ethics, and specimens

The study combined knowledge-guided analyses of public single-cell RNA-sequencing datasets with a TNBC study cohort and paired tissue metabolomics. The TNBC study cohort comprised seven patients. Each patient contributed a biopsy and surgery specimen, and patient P3 contributed an additional recurrence specimen, giving 15 single-cell specimens. The metabolomics analysis included the 14 paired biopsy and surgery specimens; no recurrence metabolomics specimen was available. Each metabolomics specimen was measured in three technical injections. Positive- and negative-polarity spectra were acquired within the same injections and separated computationally, and technical injections were not treated as biological replicates.

Patient identifiers P1–P7 were used throughout the figures and analysis files. The retained clinical grouping classified P1, P5 and P7 as non-metastatic and P2, P3, P4 and P6 as metastatic. Metastatic status was distinct from sampling time point and from the response and T-cell clonotype-expansion strata analyzed in the public cohorts. All patients provided written informed consent in accordance with the Declaration of Helsinki. The study was approved by the Ethics Committee of Tata Memorial Centre, Mumbai (TMC-IEC III; DCGI registration number IEC III: ECR/149/Inst/MH/2013/RR-24; approval reference number 900966/2023). Clinicopathological characteristics are reported in Supplemental Table 4A.

### Method details

#### Study design and overview of the scKNIFE framework

scKNIFE integrated a non-negative gene-by-cell expression matrix with a heterogeneous biological knowledge graph. The graph connected measured genes to pathways, cell identities, signaling components, metabolic reactions, and metabolites. Graph-regularized non-negative matrix factorization then learned node-factor loadings and matched cell-factor activities for each dataset independently.

The fitted matrices provided a shared representation for observed genes and graph-linked biological entities. Reconstructed node activities were used for latent-space clustering, cell annotation, differential activity analysis, metabolic-task and pathway scoring, and activity-based signaling analysis. These quantities were interpreted as model-derived relative activities rather than direct measurements of pathway flux, metabolite concentration, or molecular communication.

The study cohort was also characterized through an independent scRNA-seq workflow comprising scVI integration, community detection, multi-source cell-type annotation, transcriptome-derived copy-number inference, and cell-level TNBC subtype scoring. The independent study-cohort and scKNIFE workflows shared aligned retained cells and raw counts but used separate preprocessing and representations. scVI coordinates, communities, and annotations were not used to construct or fit scKNIFE.

#### Construction of the biological knowledge graph

scKNIFE represented prior biological knowledge as an undirected graph whose nodes included genes and annotated functional entities. The fixed benchmark graph integrated task–reaction and reaction–gene mappings from the human scCellFie resource, CellChat interaction entities, MSigDB Hallmark gene sets, cell-type marker sets, cell-state gene sets and the Enrichr Gene Ontology Biological Process 2021 collection ^8,9,14,47,48^. Cell-type priors combined human PanglaoDB markers, manually curated canonical markers, TNBC markers and CellTypist-derived marker information ^44,46^. Individual source graphs were merged through shared node identifiers. Repeated unordered edges were collapsed in the simple fitting graph, while source and relation metadata were retained as attributes. CellChat pathway names were retained as interaction annotations and were not counted as fitted latent factors.

The benchmark graph contained 22,951 nodes and 216,079 edges: 14,525 genes, 6,036 Gene Ontology terms, 1,157 CellChat entities, 787 reactions, 218 metabolic tasks, 95 cell types, 83 cell states and 50 Hallmark gene sets. The full graph used for analyses of the TNBC study cohort added Human-GEM/Human1 genes, reactions, metabolites and pathways ^13^. It contained 39,247 nodes and 336,225 edges: 14,665 genes, 12,666 reactions, 6,036 Gene Ontology terms, 4,130 metabolites, 1,157 CellChat entities, 218 metabolic tasks, 147 Human-GEM pathways, 95 cell types, 83 cell states and 50 Hallmark gene sets. Both graphs contained 33 connected components. The reported coverage totals of 7,521 and 7,668 summed configured annotation-category memberships; they were not deduplicated biological mechanisms or fitted factor counts.

Human-GEM SBML records supplied gene–reaction, reaction–metabolite and pathway–reaction relationships, with gene–metabolite shortcut edges added through shared reaction neighborhoods. Metabolites were merged by sanitized display names across model compartments while compartment identifiers were retained as attributes. The fitting graph was undirected and did not retain reaction direction or stoichiometry as a mass-balance constraint. Consequently, metabolite, reaction and pathway outputs were transcriptome-informed activities rather than compartment-resolved concentrations or metabolic fluxes. Complete resource harmonization and graph-topology procedures are provided in Methods S1 and Supplemental Table 2.

#### Expression preprocessing and gene-to-graph alignment

Each dataset was processed independently. Genes were ranked by mean expression, and the leading 7,000 genes were selected before graph intersection. For raw count *Y*_*gc*_ for gene *g* in cell *c*, counts per million (CPM) and the transformed expression value were

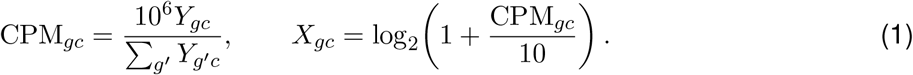

The resulting non-negative matrix was oriented as 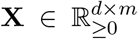, with genes in rows and cells in columns. Genes absent from the applicable knowledge graph were removed after transformation. Final fitted expression dimensions were 6,034 genes for Bassez cohort 1, 6,064 for Bassez cohort 2, 6,073 for Shiao, 5,873 for Wu TNBC, 6,047 for Tahoe, and 6,289 for the all-cell model for the TNBC study cohort. The epithelial model for the TNBC study cohort used the corresponding graph-aligned expression rows for its 43,574 selected epithelial cells. This transformation was distinct from the library-size normalization and natural-log transformation used for expression-based annotation and marker displays.

Expression-gene symbols were matched case-insensitively to graph gene nodes. The binary mapping matrix **R**_*g*_ *∈ {*0, 1*}*^*d×n*^ was defined by

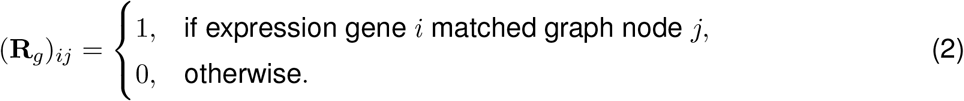

The mapping was constructed separately for each dataset while the corresponding benchmark or full graph topology was held fixed.

#### Graph-regularized non-negative matrix factorization

scKNIFE extends non-negative matrix factorization so that latent factors are defined over nodes in a biological knowledge graph rather than over measured genes alone. The factors remain anchored to the observed single-cell expression matrix while assigning loadings to connected entities such as pathways, cell states, reactions, metabolites and signaling components. This provides paired representations of biological-entity composition and factor activity across cells.

Let 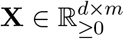 denote the non-negative gene-by-cell expression matrix, where *d* is the number of retained genes and *m* is the number of cells. Let **R**_*g*_ *∈ {*0, 1*}* ^*d×n*^ map these genes to the *n* graph nodes. The graph was represented by a binary adjacency matrix **A**, diagonal degree matrix **D**, and unnormalized Laplacian **L** = **D** *−* **A**. The fitting graph therefore had a numerically non-negative adjacency matrix. This representation records the presence of a curated relationship; it does not assign an activating or inhibitory sign, causal direction or biological effect magnitude to that relationship.

The model learns a non-negative node-factor loading matrix 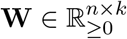 and cell-factor activity matrix 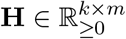 by minimizing

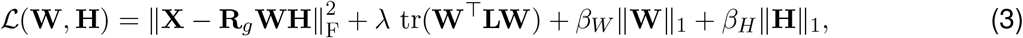

subject to **W** *≥* 0 and **H** *≥* 0. Here, *W*_*ik*_ is the loading of graph node *i* on node factor *k*, and *H*_*kj*_ is the activity of the matched cell factor *k* in cell *j*. The reconstructed activity of graph node *i* in cell *j* is (**WH**)_*ij*_.

The reconstruction term anchors the paired node and cell factors to observed gene expression. The graph term encourages connected nodes to receive similar node-factor loadings because

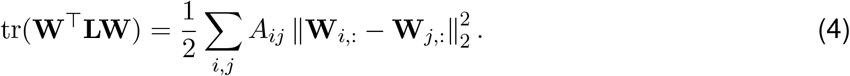

The *l*_1_ penalties controlled sparsity in the node-factor loadings and cell-factor activities. The coefficient *λ* controls graph smoothing: larger values place greater emphasis on similarity between connected node loadings. The coefficients *β*_*W*_ and *β*_*H*_ control sparsity in the node-factor and cell-factor representations, respectively. Larger values generally produce more selective representations, whereas smaller values permit broader factor composition or more mixed factor activity. The latent rank *k* and all three regularization coefficients were selected separately for each dataset; production settings are reported in Supplemental Table 3A.

#### Model initialization and multiplicative-update optimization

Only graph nodes matched to expression genes were directly observed. Standard non-negative matrix factorization with NNDSVDa initialization was fitted to the uniquely mapped gene submatrix for at most 1,000 iterations. The resulting gene loadings were copied into the corresponding rows of the initial node-factor loading matrix **W**^(0)^. The initial cell-factor activity matrix **H**^(0)^ was sampled from the uniform distribution on [0, 1).

The unobserved rows of **W**^(0)^ were initialized by solving

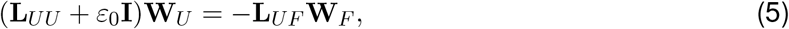

where *F* and *U* denote the observed and unobserved graph-node sets, respectively, and *ε*_0_ = 10^*−*6^ was used for numerical stability. The system was solved column-wise by conjugate gradient, and negative numerical residuals were truncated to zero.

This initialization separates directly expression-anchored gene nodes from graph-only entities. Harmonic extension supplies graph-only nodes with a graph-consistent starting value, but does not imply that those entities were directly measured.

For compact notation, define

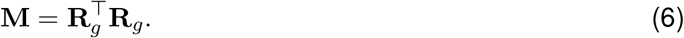

The gradients of Equation (3) are

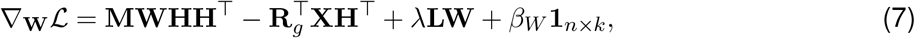

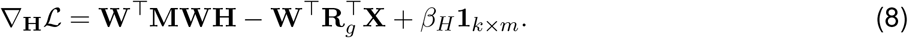

Separating each gradient into non-negative positive and negative terms gives the alternating updates

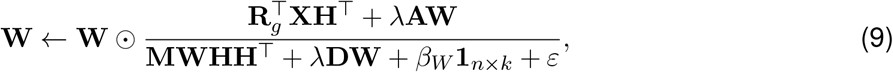

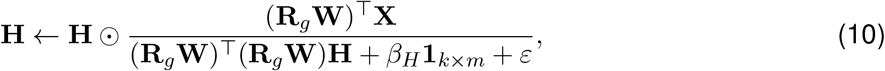

where *⊙* and the displayed fractions denote elementwise multiplication and division, standard matrix products retain their usual meaning, and *ε* = 10^*−*9^ prevents division by zero. Because every numerator and denominator is non-negative, the updates preserve **W, H** *≥* 0^49^. Each dataset was fitted independently for at most 500 iterations. The fitted matrices **W** and **H** contained node-factor loadings and cell-factor activities, respectively. Reconstructed node activities were calculated as **A**_node_ = **WH**.

Thus, the reconstruction and graph terms in the monitored value differ by a factor of two from the update objective in Equation (3); saved *β* values are reported on the implemented scale without retrospective rescaling. Where enabled, early stopping required relative change in the monitored normalized loss to remain below the configured tolerance for ten consecutive iterations; tolerance zero disabled this rule. Initial NMF used random_state=42, whereas the historical joint optimizer drew **H**^(0)^ from NumPy’s global random state, so deterministic seeding of every production fit was not assumed. The all-cell and epithelial fits for the TNBC study cohort ran for 500 iterations with early stopping disabled.

#### Projection of cells excluded from joint optimization

For the largest external datasets, joint optimization was performed on a retained within-dataset cell subset and the remaining cells from that same dataset were projected onto the fixed fitted basis. This procedure reduced the joint-optimization workload; it was not cross-dataset transfer or validation on an independently fitted cohort. Initial gene-level NMF preceded the split and used all retained cells.

Let 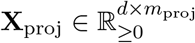 denote expression for cells reserved from joint optimization and let

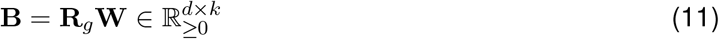

be the fixed fitted basis in expression space. For each projected cell *j*, activities were estimated by non-negative least squares,

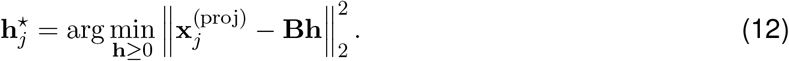

Equivalently,

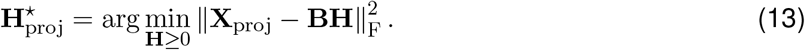

The projection did not apply the *β*_*H*_ penalty used during joint optimization. Bassez cohort 1 used 49,272 jointly optimized and 32,848 projected cells; Shiao used 49,999 and 212,071; and Tahoe used 50,000 and 176,350. Bassez cohort 2, Wu TNBC and both models for the TNBC study cohort were jointly optimized over all retained cells.

#### Implementation of benchmark latent-variable models

To place scKNIFE in the context of interpretable latent-variable frameworks, we benchmarked it against Spectra ^5^, SLALOM ^7^, and expiMap ^6^. Each comparator was fitted independently to the same cohort-specific expression data using its supported structured priors, and its cell-resolved latent representation was retained for evaluation. Comparisons used matched cells and the frozen outputs described under Quantification and Statistical Analysis. The exact prior construction, filtering thresholds, model parameters, training schedules, and embedding-extraction procedures are provided in Methods S3.

#### scKNIFE latent-space clustering

External-cohort clustering used the final cell-factor representation **H**^*⊤*^ from each graph-regularized factorization. Cell-factor columns were assembled in numerical order, reduced by principal component analysis, and partitioned using Leiden community detection on cosine-distance nearest-neighbor graphs. For the TNBC study cohort, scKNIFE was fitted before reference annotation; SingleR classified individual cells rather than H-graph clusters. The saved atlas UMAP was computed separately from the post-fit cell-factor representation **H**^*⊤*^ for visualization.

Benchmark configurations were ranked by silhouette among solutions matching the number of published author-defined cell classes. Author labels were used for post-hoc agreement evaluation but not as scKNIFE factorization targets. Bassez A was additionally evaluated with an unconstrained silhouette-selected solution. Complete tuning summaries are provided in Methods S4 and Tables S3B and S3C.

#### Earlier benchmark cluster annotation

For external-cohort benchmarks, cell types were assigned after clustering by averaging scKNIFE cell-identity and marker-node activities within each finalized Leiden community and selecting the highest-scoring available marker feature. This earlier activity-based workflow was used for benchmark evaluation, not to generate the displayed SingleR labels in the TNBC atlas.

#### Cell-level SingleR annotation of the TNBC atlas

Cell-level reference annotation was performed using SingleR v2.4.1^50^ and the full Wu breast-cancer atlas, comprising 100,064 reference cells from 26 donors ^1^. Reference and query raw counts were independently normalized to 10,000 counts per cell and natural-log1p transformed. The normalization denominator included all 29,733 reference genes and all 16,456 available query genes in their respective matrices; the normalized matrices were then restricted to 16,456 shared genes. Wilcoxon reference-marker detection and within-label reference aggregation with power 0.5 were used for training. The resulting classifier was applied separately to each of the 69,391 query-cell expression profiles, using score quantile 0.8, fine-tuning threshold 0.05, and default SingleR outlier-based pruning. Aggregation applied to the reference; each query cell retained its own prediction.

Major and minor reference labels were predicted independently. Raw predictions, pruned labels, and confidence margins were retained by cell ID; missing pruned labels were recorded as ambiguous. Figure 4 retains the original 12-category annotation alongside independently assigned cell-level full-Wu SingleR annotations. The nine major reference labels were represented as eight broad display groups by combining cancer and normal epithelial predictions into Epithelial. The minor tier retained 29 reference identities after pruning, with 72 ambiguous cells. Cell-level annotations were aligned by cell ID to the saved *H*-derived UMAP for visualization. These labels were not factorization targets. Figure 7 uses 69,319 joined cells, eight broad SingleR groups for the CellChat comparison, and 29 pruned minor identities for the resolved PDGF matrix. Supplemental Table 4C preserves the cell-level crosswalk; the broad and minor tiers are descriptive annotation layers rather than a forced hierarchy.

Canonical-marker expression was displayed as annotation context and was not individually thresholded. Cell states were assigned separately from graph-encoded state-node activities. Reference details and annotation crosswalks are provided in Methods S4 and Supplemental Table 4.

#### Metabolic evidence integration and reaction-rule comparison

Multilayer evidence-integrated activity (MEIA) combined a direct task or pathway node with its mapped reaction and gene support. The task layer used 218 human scCellFie tasks, and the pathway layer used 138 analyzable Human-GEM pathways. For entity *e* in cell *c*,

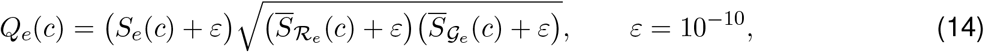

where *S*_*e*_ is direct node activity and the barred terms are mean activities over the entity’s annotated reactions and genes. Task and pathway blocks were normalized separately to sum to one within each cell. Direct scKNIFE node scores received the same block-specific normalization for comparison. A lower MEIA value therefore denoted lower relative allocation within its feature block rather than an absolute decrease in reaction flux or metabolite concentration.

A separate gene–protein–reaction-derived pathway analysis parsed Human-GEM rules independently in every cell. AND relationships used the minimum component-gene activity, and OR relationships used the maximum among alternative genes or isozymes. Reaction values were divided by their reaction-specific 99th percentiles and clipped to [0, 1]. GPR pathway activity was the mean of its mapped reaction values and was expressed relative to the mean of the other 137 pathway activities:

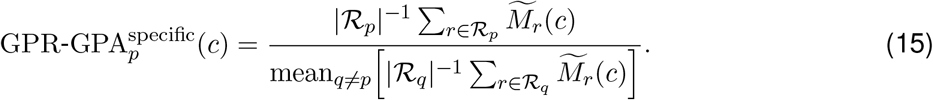

This construction used gene-rule support and was compared with, rather than multiplied by, direct scKNIFE pathway activity.

Entity-wise Pearson and Spearman correlations between MEIA and direct normalized scores used 69,390 shared cells and 356 matched entities. Within-cell correlations compared the corresponding 356-entry profiles and were summarized by patient. GPR-GPA-specific scores were compared with direct scKNIFE scores over the same 138 pathways. These analyses assessed technical consistency among related transcriptome-derived representations and did not establish accuracy against measured pathway flux. Complete mapping and screening procedures are provided in Methods S5 and S6.

#### Cross-cohort node-factor concordance and within-cohort redundancy

We compared the final node-factor loading matrices **W** from Bassez cohorts A and B, Shiao and Wu. Each matrix contained one row for each of 22,951 graph nodes and one column for each fitted latent factor. Graph-node identifiers recovered from the corresponding fitted objects were required to match the **W** row count and to occur in an identical sequence across all four datasets.

For cross-dataset comparison, each **W** column was standardized to zero mean and unit variance. The standardized matrix was divided by the square root of its number of factors, each graph-node row was normalized to unit Euclidean norm, and the resulting matrix was column-centered. For two processed matrices **U** and **V**, linear centered-kernel alignment (CKA) was calculated as ^51^

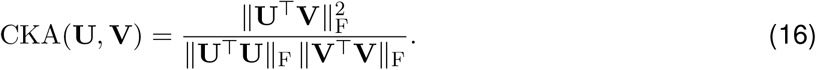

This statistic compares the graph-node geometries represented by **W** without requiring equal factor numbers or one-to-one matching of individual factors.

Within each dataset, factor redundancy was evaluated from the column-standardized **W** matrix. Pairwise dependence was summarized by the median, 95th percentile and maximum of the absolute Pearson correlations between distinct factor columns. Effective rank was calculated from the singular values *s*_*i*_ of standardized **W** ^52^. With normalized spectral weights

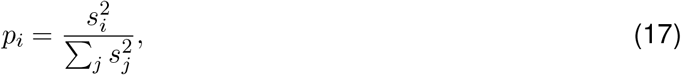

the effective rank was exp(*− ∑*_*i*_ *p*_*i*_ log *p*_*i*_), and the effective-rank fraction was obtained by dividing this value by the fitted factor count. The standardized-matrix condition number was calculated as the ratio of its largest to smallest singular value. These quantities were treated as descriptive diagnostics of representational concordance and redundancy; no null-hypothesis test or *P* value was assigned.

For pooled visualization, the four independently processed **W** blocks were concatenated by graph-node row, producing a 22,951-by-190 matrix, and rows were normalized again to unit Euclidean norm. A t-distributed stochastic neighbor embedding (t-SNE) was calculated from this pooled representation ^53^. Separate t-SNE coordinates were also calculated for each independently normalized cohort block. Every map contained the same graph nodes in the validated common order; coordinates were not used for model fitting or quantitative distance comparisons.

Communities were defined from the pooled high-dimensional representation rather than from the two-dimensional t-SNE coordinates. A cosine-neighbor graph was partitioned by Leiden community detection ^54^. A resolution sweep from 0.25 to 2.0 yielded 5–34 communities. Resolution 1.0 was retained for display and produced 18 communities; these pooled labels were transferred unchanged to the pooled and cohort-specific maps rather than re-estimated within each cohort. Community size, modularity and node-type composition were calculated from the frozen pooled assignments. The maps and community summaries were descriptive and were not treated as annotation-enrichment tests or evidence of independently recovered cohort-specific partitions.

#### Graph-node organization of shared scKNIFE node factors

Each latent dimension paired a node-factor loading profile in **W** with its matched cell-factor activity profile in **H**. The epithelial program analysis evaluated 50 aligned node factors by clustering their graph-node profiles from **W** after excluding cell-type and cell-state annotation nodes and zero-signal rows. The current analysis retained nine W-derived node groups (NG1–NG9). Twenty globally unique representative nodes were selected per group by ranking positive node enrichment with exponential weights exp[*−*(*r −* 1)*/*5] and retaining ranks through 95% cumulative weight, subject to a maximum of 20 nodes per group. The resulting 180-node display summarizes group composition; group labels are descriptive and do not imply independently validated pathways. This unsupervised organization was not a null-hypothesis test.

The matching **H** rows provided cell-resolved activities for the programs defined from **W**. Selected node scores were reconstructed as max(**HW**_node_, 0), transformed with ln(1 + *x*), and standardized across the epithelial cohort for display. P4/P1 matched-sample comparisons used distinct biopsy and surgery cells after cell-type and RNA-depth matching; they are descriptive within-patient summaries. Equal-patient and cell-weighted summaries were retained separately because patients contributed different cell numbers. Supporting patient and estimator summaries are provided in Methods S7 and Supplemental Table 5.

#### Activity-based cell–cell communication scoring

Because single-cell RNA sequencing does not directly observe communication events, exploratory communication scores were calculated from reconstructed scKNIFE node activities for ligands, receptors and annotated modulators. These activities are model-conditioned estimates and should not be interpreted as direct measurements of ligand secretion, receptor protein abundance, receptor occupancy or downstream signaling.

CellChatDB Secreted Signaling, ECM–Receptor, and Cell–Cell Contact interactions were retained, with receptor complexes expanded into component records. The activity-based communication score is distinct from CellChat communication probabilities and does not estimate treatment-induced changes in communication.

The CellChat comparison used the 69,319 joined atlas cells assigned to eight broad SingleR groups. The resolved PDGF sender–receiver matrix used the 29 independently predicted, pruned minor identities; these minor labels were not treated as a nested extension of the broad groups.

For each source cell type *s* and target cell type *t*, we first summarized ligand and receptor activity at the cell-type level. For a ligand *l* and receptor *r*, we defined

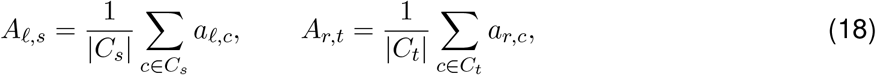

where *a*_*g,c*_ denotes the scKNIFE activity score of ligand or receptor *g* in cell *c*. The sets *C*_*s*_ and *C*_*t*_ contain cells assigned to the source and target cell types, respectively. Thus, *A*_*l,s*_ measures ligand activity in the sender population, whereas *A*_*r,t*_ measures receptor activity in the receiver population.

To favor interactions selective for the relevant cellular context, specificity for ligand or receptor *g* in cell type *u* was defined as

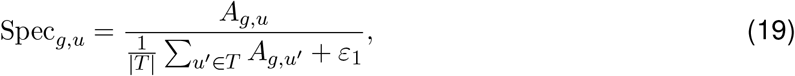

where *T* is the set of cell types and *ε*_1_ = 10^*−*10^. Pair specificity was Spec_*lr*_ (*s, t*) = Spec_*l,s*_Spec_*r,t*_. Each ligand–receptor pair also received an evidence weight *C*_*lr*_ and an activity-supported cofactor multiplier Ω_*lr*_(*s, t*).

Combining these components, the communication contribution of ligand *l* and receptor *r* from source cell type *s* to target cell type *t* was defined as

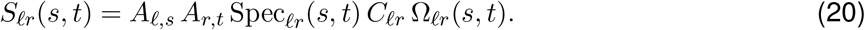

This formulation assigns a high score when the ligand is active in the sender, the receptor is active in the receiver, the interaction is cell-type selective, and its evidence and cofactor context are favorable. The resulting score quantifies compatibility between cohort-level sender and receiver activity profiles. It does not demonstrate physical contact, signaling in individual cells or causal pathway activation. Exact evidence weights, cofactor multipliers, and network-summary definitions are provided in Methods S9.

The total communication score from source cell type *s* to target cell type *t* was obtained by summing over all valid ligand–receptor components,

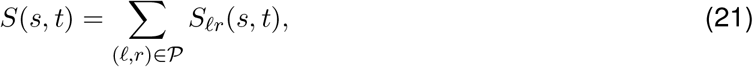

where *P* is the retained interaction set. Pathway-specific matrices were obtained by summing contributions within CellChatDB pathways, and sender and receiver strengths were calculated from matrix row and column sums.

Evidence weights were 1 for KEGG-supported records, 0.8 for other documented evidence, and 0.5 when evidence was missing. Positive reconstructed activities of annotated agonists, antagonists, activating co-receptors, and inhibitory co-receptors applied heuristic multipliers of 1.5, 0.5, 1.3, and 0.7, respectively. These values were analysis weights rather than measured receptor occupancy or biochemical rate constants.

#### Tissue preparation, Flex libraries, and initial single-cell processing

All 15 tissues used for single-cell profiling were received fresh-frozen and stored at *−*80 °C until processing. Tissue fixation and dissociation were performed according to 10x Genomics protocol CG000553, revision B, without modification. Approximately 25 mg of minced tissue was resuspended in freshly prepared fixation buffer containing 4% formaldehyde in 1*×* Fix & Perm buffer and incubated for 18 h at 4 °C. Fixed tissue was dissociated at room temperature using a tissue dissociator and pre-warmed RPMI-1640 containing 2 mg mL^*−*1^ Liberase TL. Human transcriptome probe pairs were then hybridized and ligated, followed by quenching and washing. Cell counts and viability were assessed by Ethidium Homodimer-1 exclusion.

Multiplexed gene-expression libraries were prepared following 10x Genomics protocol CG000527 without modification. Each of four libraries contained four probe-barcode assignments, BC001–BC004; these sample identifiers were distinct from the i7/i5 sequencing indices. GEM generation was performed on a Chromium X instrument (10x Genomics). Final library size distributions and concentrations were assessed on an Agilent TapeStation using the High Sensitivity D5000 ScreenTape assay, and libraries meeting the expected fragment-size and molar-concentration criteria were sequenced. Libraries were indexed using Dual Index Kit TS, Set A (PN-1000251). SI-TS-A1, SI-TS-B1, SI-TS-C1, and SI-TS-D1 were assigned to libraries 1–4, respectively.

Libraries were sequenced on an Illumina NovaSeq X Plus using a 1.5B flow cell and paired-end 2 *×* 150-bp reads. The i7 and i5 index reads followed the configuration recommended by 10x Genomics. The sequencing design targeted approximately 10,000 read pairs per cell and 10,000 cells per library–sample slot, corresponding to approximately 1.6 billion read pairs over the 16 multiplexed slots. Demultiplexed FASTQs were processed with Cell Ranger 9.0.0 cellranger multi, using refdata-gex-GRCh38-2024-A and Chromium_Human_Transcriptome_Probe_Set_v1.1. 0_GRCh38-2024-A.csv. The four configurations specified Gene Expression libraries, sample-specific probe barcodes, and create-bam=true; Flex counts were assigned by probe-pair alignment and UMI aggregation, and per-sample filtered feature–barcode matrices supplied the downstream cells.

The retained lineage used library1_new for library 1 and libraries 2–4, comprising 16 library–sample entries and 80,217 Cell Ranger-called cells before downstream filtering.

#### Tissue metabolite extraction and LC–MS acquisition

Tissue was stored at *−*80 °C and pulverized under liquid nitrogen. Approximately 30 *±* 5 mg was extracted with cold methanol–water (4:1, v/v) at 150 *µ*L per 10 mg tissue, using glass or ceramic beads. Homogenization used three 20-s cycles at 6,000 rpm separated by 5 s, with dry-ice cooling. Samples were incubated for 1 h at *−*20 °C and centrifuged at 16,000 *× g* for 15 min at 4 °C. Supernatants were dried in a vacuum concentrator at 10 °C, reconstituted in methanol–water (1:1, v/v) with volume normalized to tissue mass, sonicated for 1 min in an ice-cold bath, centrifuged again at 16,000 *× g* for 15 min at 4 °C, and transferred to LC vials.

Reversed-phase separation used a Waters BEH C18 column (2.1 *×* 100 mm, 1.7 *µ*m particles, 130-Å pores) at 400 *µ*L min^*−*1^. Mobile phase A was water with 0.1% formic acid and B was acetonitrile with 0.1% formic acid. The programmed B proportion was 1% at 0 and 1 min, increased to 99% at 10 min, remained at 99% through 13 min, returned to 1% at 14 min, and remained at 1% through 17 min.

Acquisition used a Thermo Scientific Orbitrap Exploris 120 with electrospray ionization. Retained metadata recorded Xcalibur 4.3 builds 4.3.373.8 and 4.3.458.15. Positive- and negative-polarity full scans and data-dependent tandem scans were acquired within the same RAW files. Inspected acquisition metadata recorded full scans over *m/z* 100–1,000 at resolving power 120,000 and tandem scans at resolving power 60,000; these recorded settings were used in preference to the protocol narrative’s generic *m/z* 50–1,000 range. The retained protocol described a nominal 10–50-eV collision-energy range, but the complete run-wide collision-energy and scan-timing program requires the instrument-method export.

#### RAW conversion, polarity separation, and LC–MS preprocessing

ProteoWizard converted 42 sample acquisitions and three solvent blanks from 14 biopsy/surgery specimens (three injections each) to centroided mzML; MSnbase and mzR separated positive and negative modes. OptiLCMS and MetaboAnalystR processed each mode separately with blank subtraction, contaminant removal, peak detection, alignment, and isotope/adduct annotation. Technical injections were summarized per specimen, not treated as independent samples. No pooled quality-control samples were available for drift correction.

#### Study-cohort scRNA-seq preprocessing and quality control

The study-cohort count matrix was processed independently of scKNIFE. Quality control excluded cells with fewer than 200 detected genes, more than 20% mitochondrial counts, or a stored doublet call of 1, and genes detected in fewer than three cells. The retained analysis object contained 69,391 cells and 16,456 genes; quality-control diagnostics are provided in Methods S11.

#### Integration, clustering, and cell-type annotation

The independent analysis selected 3,000 highly variable genes using the batch-aware Seurat-v3 procedure, with library as the batch key. scVI ^41,42^ was fitted to the count matrix with library as the batch annotation and produced a ten-dimensional representation. Leiden clustering on the scVI neighbor graph yielded 21 communities, selected using silhouette coefficients and stability across adjacent resolutions. Harmony ^43^ was evaluated as a comparison method but was not used for clustering.

The 21 communities were annotated using canonical markers, CellTypist predictions ^44^, PanglaoDB enrichment ^46^, scArches mapping to the Wu reference ^45^, published breast and TNBC markers, and expression-module scores, yielding 12 cell types. The scVI coordinates, communities, and annotations were not used to construct, tune, initialize, or fit scKNIFE. Annotation and validation details are provided in Methods S11.

#### Marker identification

Marker genes were ranked using the Scanpy Wilcoxon procedure, with individual cells as observations, and used to characterize cell types. Transcriptome-derived CNV inference and TNBC subtype scoring are described in Methods S11.

### Quantification and statistical analysis

#### Statistical framework and experimental units

Patients were the experimental units for patient-level analyses, with repeated specimens paired within patient. Cell-level analyses were exploratory. Tests were two-sided unless noted; Holm adjustment was used for LC–MS pathway Gamma probabilities, and Benjamini–Hochberg correction ^55^ for the screening families below. Selected node comparisons are reported with nominal post-selection *P* -values. Effect sizes, confidence intervals, and directional concordance were considered alongside *P* -values; statistical significance alone was not taken as evidence of causality.

Downstream analyses used frozen, checksummed outputs and did not rerun model fitting, differential models, or UMAP optimization unless stated otherwise. Seeds are reported when retained run records identify them. UMAPs, heatmaps, overlap diagrams, and network layouts are descriptive unless a formal test is specified. Patient sample sizes accompany patient-level comparisons; adjusted *P* -values are denoted as *q* where useful, and values below machine precision are not reported as zero. Selected examples and cell-level screens are interpreted as exploratory.

#### Software and reproducibility

Initial single-cell processing used Cell Ranger 9.0.0. The CNV-analysis branch used Python 3.9.21, Scanpy 1.9.6, and infercnvpy 0.4.5; LC–MS conversion and polarity separation used ProteoWizard 3.0.26068, MSnbase 2.32.0, and mzR 2.40.0. Saved metabolomics sessions used R 4.4.3, limma 3.62.1, OptiLCMS 1.2.0, and MetaboAnalystR 4.2.0. These versions correspond to individual analysis components; additional software details are listed in the Key Resources Table.

Analysis scripts used frozen model outputs, metadata, statistical tables, and documented transformations. Supplemental Tables 1–7 provide graph inventories, parameter-selection records, annotation crosswalks, program definitions, and statistical summaries.

#### Cell-type-aware differential activity analysis

Differential analyses used the cell-by-node activity matrix produced by scKNIFE. Comparisons were performed within each assigned cell type to limit effects caused by changing cell-type abundance. For every prespecified contrast, node activities were compared between reference and comparison groups using a two-sided Wilcoxon rank-sum test, equivalently expressed as the Mann–Whitney *U* test.

Reported summaries included group sizes, means, medians, test statistics, log_2_ mean fold changes, and effect sizes. Benjamini–Hochberg correction was applied over the hypothesis family defined by each analysis workflow. Analysis-specific correction families, thresholds, and interpretation boundaries are reported in the corresponding Quantification and Statistical Analysis or Supplemental Methods subsection.

#### Evaluation of computational benchmarks

Clustering agreement was assessed using adjusted Rand index, adjusted mutual information, and normalized mutual information on cells shared by scKNIFE, each comparator, and the reference annotations. Intersections contained 81,142, 36,264, and 38,251 cells for Bassez cohorts 1 and 2 and Wu TNBC, respectively; Shiao was omitted because matched outputs were unavailable for all comparators. These metrics were descriptive.

In matched Tahoe cells, Pearson and Spearman correlations compared MAGIC-imputed *TP53* expression with apoptosis activity and *PIK3R1* expression with PI3K–AKT–mTOR activity, using scKNIFE and Scanpy gene-set scores (64,064 and 60,743 cells, respectively). These correlations were descriptive.

#### Patient-level and exploratory cell-level metabolic analyses

Baseline scKNIFE MEIA scores were summarized as patient-level medians by cell type, requiring at least 20 cells per patient–cell-type combination and at least four Shiao or two Bassez patients per group. Group effects used Cliff’s delta, exact two-sided permutation tests of median differences, and 95% bootstrap confidence intervals (5,000 within-group resamples); Benjamini–Hochberg correction was applied separately to the Shiao response and Bassez expansion analyses.

Longitudinal task changes were summarized per patient and compared between outcome groups using exact two-sided Mann–Whitney *U* tests on the treatment-minus-baseline log_2_ median-score scale. The primary task analysis covered 218 scKNIFE MEIA tasks, with integrated and mechanistic pathways as sensitivity analyses; selected contrasts were not a separate multiplicity-controlled family. Secondary cell-level tests used two-sided Wilcoxon rank-sum or Mann–Whitney *U* tests. Bassez Gene Ontology analyses compared On and Pre cells within cohort, clonotype-expansion group, and cell type (at least 20 cells per time point), using mean log_2_ fold changes and Cohen’s *d*; Benjamini–Hochberg correction covered Gene Ontology-term-by-cell-type tests within each cohort and expansion group. These cell-level screens were descriptive; tested families are detailed in Methods S10.

#### Patient-level analysis of inferred metabolic activities

Inferred scores were summarized by patient, time point, and cell compartment. Paired biopsy– surgery differences were tested across seven patients using exact two-sided Wilcoxon signed-rank tests; recurrence values summarize the single contributing patient.

Benjamini–Hochberg correction was applied separately to node (1,344 tests), task (1,308 tests), and pathway (828 tests) screens. Node comparisons were also adjusted within each of four mapped LC–MS pathways. The focused carnitine screen covered ten nodes across six compartments (60 comparisons), with Benjamini–Hochberg correction. These transcriptome-derived scores provide relative activity estimates; patient-level summaries are in Supplemental Tables 6C–6G.

#### Permutation testing and CellChat concordance

Pooled scKNIFE communication tests permuted cell labels while holding specificity weights fixed. One-sided empirical *P* -values were calculated as (*b* +1)*/*(*B* +1) from 1,000 permutations and adjusted by Benjamini–Hochberg across 252 pathways and 2,261 ligand–receptor keys. These tests permuted cells, not patients.

Metastatic-status and sampling-site contrasts were computed within the relevant cell subsets; surgery minus biopsy defined the direction, and recurrence cells were excluded.

CellChat and scKNIFE outputs were harmonized by pathway, ligand–receptor identifier, and broad cell type. Overlap was summarized with set counts and Jaccard indices, and matched positive scores were compared on the log_10_ scale using Pearson and Spearman correlations. CellChat probabilities used saved Seurat-normalized expression, computeCommunProb with truncatedMean and trim=0.1, and at least ten cells per population. Patient-paired biopsy–surgery analyses required at least ten sender and receiver cells at both time points; exact paired sign-flip *P* -values were adjusted across 2,080 pathway/source/target comparisons.

#### Specimen-level LC–MS statistics

Non-finite and non-positive intensities were treated as missing. Technical injections were averaged on the raw-intensity scale within specimens; features present in at least 10 of 14 specimens were retained, remaining missing values were imputed as one fifth of the feature’s minimum positive intensity, and values were log_2_ transformed. Per-polarity linear models included metastatic status and sampling time point, with patient as a blocking factor, followed by empirical-Bayes moderation. Benjamini–Hochberg correction was applied across both polarities within each contrast.

#### Native MS/MS annotation and Mummichog pathway analysis

Consensus data-dependent tandem spectra were searched against MS2ID biology, lipid, pathway, and exposomics libraries. Candidate annotations were matched to features by polarity, mass, and retention time and treated as putative in the absence of authentic standards. Mummichog 2 with the human MFN model ^56^ analyzed ranked features from both polarities using mixed ion mode, 15-ppm mass tolerance, the leading 10% of features, and 100 permutations per contrast. Permutation-background Gamma probabilities were adjusted using Holm correction.

## RESOURCE AVAILABILITY

### Lead contact

Requests for further information and resources should be directed to and will be fulfilled by the lead contact, Debarka Sengupta.

### Materials availability

This study did not generate new unique reagents.

### Data and code availability

The raw LC–MS data, converted mzML files, processed feature-intensity matrices, putative metabolite annotations and associated sample metadata have been deposited in MassIVE under accession MSV000103682 (doi:10.25345/C5833ND1W).

The longitudinal TNBC single-cell dataset generated in this study is available from the European Nucleotide Archive under study accession PRJEB108811 (secondary accession ERP189646; public release date April 30, 2026). All other datasets analyzed in this study are publicly available from their original sources. Supplemental Tables 1 through 7 are provided separately as tab-delimited files and consolidated Excel workbooks. The Bassez A and B breast-cancer single-cell RNA-sequencing cohorts, the TNBC subset of the breast-cancer atlas from Wu et al., the longitudinal TNBC dataset from the Shiao et al. clinical-trial resource, and plate 7 of Tahoe-100M (including doxorubicin- and afatinib-treated cell lines) are described in Methods S2 (“Public datasets, model selection, and large-dataset projection”). The independent annotation benchmark used lung-cancer profiles from the Dong et al. resource, retrieved through CELLxGENE. The scKNIFE source code is available at https://github.com/bernadettem03/scKNIFE. Processed data and additional information needed to reanalyze the results will be provided by the lead contact upon reasonable request.

## ACKNOWLEDGMENTS

D.S. acknowledges support from the iHub-Anubhuti-IIITD Foundation under the NM-ICPS scheme of the Department of Science and Technology (DST), the Science and Engineering Research Board (SERB grant CRG/2022/007706), the Department of Biotechnology (DBT grant IC-12044(12)/4/2022-ICD-DBT), and the Indian Council of Medical Research (ICMR grant IIRPIG-2024-01-01321).

## AUTHOR CONTRIBUTIONS

Conceptualization, D.S.; experimental design, S.H., B.M., and D.S., with support from S.J. and S.G.; investigation, B.M., Akash Maity, and N.G.; mathematical methodology, D.S., with support from Angshul Majumdar, B.M., A.H., M.F., S.S., and S.K.; software and formal analysis, B.M., with support from R.G.; validation and visualization, S.P., J.T.S., and N.B.; supervision, D.S.; writing—original draft and writing—review and editing, all authors. All authors reviewed and approved the manuscript.

## DECLARATION OF INTERESTS

The authors declare no competing interests.

## DECLARATION OF GENERATIVE AI AND AI-ASSISTED TECHNOLOGIES

During the preparation of this work, the authors used generative AI tools to assist with language editing, improving clarity, and code debugging. All AI-assisted text and code were reviewed, edited, and validated by the authors. The authors take full responsibility for the accuracy, integrity, and conclusions of the work.

## SUPPLEMENTAL INFORMATION INDEX

Supplemental Results extend the cross-cohort representation analysis, benchmark comparisons, external-cohort metabolic candidates, study-cohort annotation, epithelial programs, tissue-metabolomics integration and activity-based communication reported in Figures 2–7. Supplemental Methods S1–S11 describe graph provenance, datasets and model selection, comparator configurations, clustering and annotation, metabolic reconstruction and screening, epithelial node-group construction, patient-resolved activity summaries, communication analyses, longitudinal Gene Ontology analysis and independent processing of the study-cohort single-cell data. Supplementary Figures S1–S2, S3A–S3B and S4–S6 provide the corresponding extended visualizations. Supplemental Tables 1A–1B, 2, 3A–3D, 4A–4F, 5A–5G, 6A–6I and 7A–7G provide graph inventories, model-selection records, cohort annotations, epithelial-program definitions, metabolic summaries and communication results; Supplemental Tables 7A–7G support main Figure 7. Supplemental Results, Methods, table legends and figure legends are provided in the Supplemental Information; tab-delimited data tables and consolidated Excel workbooks are supplied separately.

## Notes

### Competing Interest Statement

The authors have declared no competing interest.

### Summary of Updates

This version clarifies how scKNIFE connects cellular organization, biological programs, metabolism and cell-cell communication within a shared framework. The Results and Discussion have been revised to strengthen the biological interpretation, particularly the links between tissue metabolomics and cell-type-specific metabolic activities. Figures, legends and supplementary tables have been updated for consistency. The Methods and statistical reporting have been clarified, and the data and code availability sections now include repository details and accession numbers.

