## Supplementary File for "A knowledge graph-guided framework for multitask single-cell analysis"

### SUPPLEMENTAL RESULTS

#### Cross-cohort node representations retain shared structure and cohort-dependent variation

Graph-node loading matrices $\mathbf{W}$ from Bassez cohorts 1 and 2, Shiao, and Wu TNBC shared 22,951 ordered nodes and contained 40, 40, 80, and 30 factors, respectively. Centered-kernel alignment ranged from 0.669 to 0.895, highest between the Bassez cohorts (0.895), while effective ranks ranged from 15.78 to 56.80 (Supplementary Figure S1; Supplemental Table 1A)[1].

The pooled representation comprised 190 factor features across the shared nodes. A single Leiden partition identified 18 communities, which were transferred to independent t-SNE views colored by graph-node kind (Supplementary Figure S1b,c; Supplemental Table 1B)[2,3].

#### Graph coverage and comparator embeddings contextualize the benchmark

The fixed benchmarking graph contained 22,951 nodes and 216,079 edges; the expanded graph contained 39,247 nodes and 336,225 edges (Supplemental Table 2). The benchmark graph included 14,525 genes, 6,036 Gene Ontology terms, 1,157 CellChat entities, 787 reactions, 218 metabolic tasks, 95 cell-type annotations, 83 cell-state annotations and 50 Hallmark gene sets. Configured category memberships summed to 7,521; the expanded graph added 147 Human-GEM pathway memberships, for 7,668. These counts describe graph size and configured annotation coverage, not unique independent mechanisms or fitted factors.

Production models used 40, 40, 30, 80 and 60 factors for Bassez cohort 1, cohort 2, Wu, Shiao and Tahoe, respectively (Supplemental Tables 3A–3C). Supplementary Figure S2 places comparator Leiden clusters and author labels on identical coordinates, providing qualitative context for the cluster-to-label agreement reported in Figure 2l.

Cells outside joint optimization in Bassez 1, Shiao and Tahoe were projected by non-negative least squares onto each dataset's fixed scKNIFE basis (Methods S2). The projected counts were 32,848, 212,071 and 176,350 cells, respectively. Their profiles appear in the corresponding Bassez 1 and Shiao embeddings and Tahoe correlations (Figure 2d–g,m–p).

#### External-cohort supplemental analyses provide context for selected metabolic candidates

The Bassez and Shiao analyses used distinct clinical strata: E/NE denote T-cell clonotype expansion/non-expansion in Bassez, whereas R/NR denote clinical response/non-response in Shiao[4,5]. Supplementary Figure S3A shows patient directional concordance, patient-level GO response interactions, and pooled-cell heatmaps of Bassez treatment-associated task activity. The GO forest plots use patient-level responses, whereas the heatmaps use pooled-cell effect estimates; cells in the latter are nested within patients. GO activity is not treated as enrichment of a differentially expressed gene list.

Supplementary Figure S3B localizes the Figure 3 candidate activities on frozen, cohort-specific embeddings. The mapped values are inferred task activities displayed within each cohort and time point; colors should not be compared quantitatively across maps. Supporting patient-level statistics and biological evidence are tabulated in Supplemental Table 3D. Baseline comparisons and longitudinal patient tests remain exploratory, and the two clinical stratifications are not interchangeable.

#### Nine-group epithelial summaries separate cell-weighted and patient-weighted patterns

Supplementary Figure S5 reports supporting views of the nine W-derived epithelial node groups. PG1–PG9 are the display labels for the retained NG1–NG9 analysis identifiers; they denote nine groups of 50 fitted factors. Patient coverage and cell-weighted versus equal-patient heatmaps show how inferred group patterns relate to cell abundance and patient representation. PG7 task distributions provide context for spermidine synthesis from ornithine and S-adenosyl-L-methionine synthesis, while assigned H-factor and node-group networks connect factors to epithelial/state annotations. P2 *MYC* and spermidine activities are shown at biopsy and surgery. The P4 and P1 estimator comparisons distinguish summaries across all epithelial cells from cell-type- and RNA-depth-matched populations; the direction of P4 PG8 changes between estimators. These are descriptive views of distinct sampled cells, not patient-level significance tests (Supplemental Tables 5A–5G).

### SUPPLEMENTAL METHODS

#### Methods S1. Knowledge-graph resource harmonization and topology construction

The benchmark graph was assembled from the retained scCellFie task–reaction and reaction–gene mappings, CellChatDB ligand–receptor components, MSigDB Hallmark gene sets, cell-type marker sets, cell-state gene sets, and the Enrichr Gene Ontology Biological Process 2021 collection. Cell-identity priors combined high-confidence human PanglaoDB markers, manually curated canonical epithelial, immune, stromal, and cycling markers, a TNBC-focused marker panel, and CellTypist-derived marker information. Duplicate node and cell type–marker assignments were removed.

CellChatDB interactions annotated as Secreted Signaling, ECM–Receptor, or Cell–Cell Contact were represented as directed records during source construction. The fitted source graph was subsequently simplified to an undirected graph, with pathway and evidence annotations retained as edge attributes. Metabolic-task priors used both task–reaction and reaction–gene relationships rather than treating the `Task_by_Gene` matrix as the only metabolic source.

The metabolic systems layer was constructed from the Human-GEM SBML Level 3 model[6]. Genes, reactions, metabolites, and pathways were represented as distinct node classes. Edges linked reactions to reactants and products, genes to reactions through gene–protein–reaction associations, and pathways to their constituent reactions. Gene–metabolite shortcut edges were added when genes and metabolites shared a reaction neighborhood. Human-GEM gene products were harmonized to gene symbols, and internal metabolic identifiers were mapped to human-readable names while retaining the original identifiers as node attributes.

The source graphs were merged through shared node identifiers, principally gene symbols. Duplicate unordered node pairs were collapsed, while database provenance and relation-specific attributes were retained. The fixed benchmark graph contained 22,951 nodes and 216,079 edges. The study-cohort graph preserved that base and added the Human-GEM layer, yielding 39,247 nodes and 336,225 edges. Its node classes comprised 14,665 genes, 12,666 reactions, 6,036 Gene Ontology terms, 4,130 metabolites, 1,157 CellChat entities, 218 metabolic tasks, 147 Human-GEM pathways, 95 cell types, 83 cell states and 50 Hallmark gene sets. Source and relation metadata were retained for downstream annotation even though optimization used the simplified graph.

##### Application of the fixed knowledge graph across benchmark datasets

For benchmarking, a second graph was assembled once and applied unchanged to Bassez A, Bassez B, Wu, Shiao, and Tahoe. Its seven source layers represented scCellFie task–reaction and reaction–gene membership[7], CellChatDB ligand–receptor interactions[8], MSigDB Hallmark membership[9], CellTypist markers[10], TNBC-relevant breast-tumor states from CellMarker 2.0[11], and gene–Gene Ontology associations from the Enrichr GO Biological Process 2021 collection[12,13].

Each benchmark source was converted to an undirected, unweighted NetworkX graph and merged by node identifier. Shared genes and reactions were represented once, and duplicate unordered node pairs were collapsed. Expression values, treatment or expansion labels, and author-provided cell annotations were not used to define graph topology. Before cross-resource deduplication, source-level node and edge counts were those reported in Supplemental Table 2. The integrated graph contained 22,951 unique nodes and 216,079 unique edges across 33 connected components; its largest component contained 22,906 nodes. For each dataset, the sparse adjacency and degree matrices defined the unnormalized graph Laplacian used for fitting.

### Methods S2. Public datasets, model selection, and large-dataset projection

#### Public single-cell datasets and cohort definitions

**Bassez et al.** Processed TNBC cells and author annotations were retained as two cohorts[4]. Cohort 1 (C1/A) contained 82,120 cells from 13 patients, including five clonotype-expanded (E), seven non-expanded (NE), and one patient without an assigned expansion label. The unassigned patient contributed to the embedding but not E/NE comparisons. Cohort 2 (C2/B) contained 36,721 cells from three E and three NE patients. E and NE were patient-level T-cell clonotype-expansion strata; they did not denote clinical response or expansion of every plotted cellular compartment. Pretreatment and on-treatment samples retained the supplied Pre and On labels.

**Shiao et al.** Immune and non-immune objects from GSE246613 were combined over shared genes[5]. Sixteen patients with paired sampling and cells expressing at least 200 genes yielded 262,070 cells. Sampling conditions were baseline (Base), pembrolizumab (PD1), and pembrolizumab with radiotherapy (RTPD1). The retained response grouping contained ten responders and six nonresponders. Immune R/NR and non-immune R1/R2/NR labels were harmonized by mapping R1 and R2 to R, while the three treatment time points remained distinct.

**Wu et al.** The TNBC subset of GSE176078 contained 42,512 cells from ten patients and was used for clustering and annotation benchmarking[14].

**Tahoe-100M.** The retained Tahoe subset contained 226,350 cells from 50 cancer cell lines[15]: 101,543 DMSO-treated, 60,743 afatinib-treated, and 64,064 doxorubicin-hydrochloride-treated cells. Both drugs were represented at 0.05 $\mu$M. Tahoe was a cancer-cell-line perturbation resource, not a patient cohort or a TNBC-only dataset.

#### Hyperparameter selection and production configurations

Each dataset used the 7,000-gene preprocessing and graph alignment described in STAR Methods. External-cohort tuning included an initial factor-number search, a local joint factor-number/graph-regularization search, and subsequent node- and cell-factor sparsity searches. Candidate summaries retained silhouette, reconstruction agreement, sparsity, and iteration counts. The $\beta_W$ selection sought loading sparsity between 0.35 and 0.55 and otherwise selected the closest boundary. Final production settings followed the saved configuration and were not assumed to be the maximum-silhouette candidate from every preceding grid. The configurations for benchmark and study-cohort fits are summarized in Supplemental Methods Table S1; candidate-level benchmarking records and deviations, including the retained Shiao joint-grid choice, are provided in Supplemental Tables 3A–3C.

**Supplemental Methods Table S1. Final scKNIFE configurations across benchmark and study-cohort analyses.**

| Analysis | $k$ | $\lambda$ | $\beta_W$ | $\beta_H$ | Joint cells | Projected cells |
|---|---|---|---|---|---|---|
| Bassez C1 | 40 | 0.1 | 0.01 | 0.05 | 49,272 | 32,848 |
| Bassez C2 | 40 | 0.01 | 0.001 | 0.05 | 36,721 | 0 |
| Shiao | 80 | 1 | 0.01 | 0.05 | 49,999 | 212,071 |
| Wu TNBC | 30 | 0.1 | 0.01 | 0.01 | 42,512 | 0 |
| Tahoe | 60 | 0.001 | 0.0001 | 0.01 | 50,000 | 176,350 |
| Study cohort all cells | 150 | 0.1 | 0.001 | 0.05 | 69,391 | 0 |
| Study cohort epithelial | 50 | 0.01 | 0 | 0.001 | 43,574 | 0 |

Bassez C1 used a seeded 60% within-cohort subsample for joint optimization. Shiao used a patient-, response-, and treatment-stratified target of 50,000 cells; stratum rounding produced 49,999. Tahoe used 50,000 cells. Remaining cells were projected by non-negative least squares against fixed $\mathbf{R}_g\mathbf{W}$ without the $\beta_H$ penalty. Because initial gene NMF used all retained cells before this split, projection reduced the joint-optimization workload but did not constitute wholly unseen-cell validation. The all-cell and epithelial fits for the TNBC study cohort used 500 iterations with early stopping disabled.

##### External clustering and perturbation comparisons

scKNIFE cell-factor profiles were reduced by PCA and used to construct cosine-neighbor graphs. Candidate grids evaluated 10 or 15 principal components, 5, 10, or 15 neighbors, and Leiden resolutions 0.1, 0.3, 0.5, and 1.0. For the retained author-label benchmark, solutions matching the published cell-type count were prioritized, with relaxation by one or two clusters when required; the highest silhouette among eligible solutions was selected. Author labels set the target partition count and evaluation reference but did not tune the scKNIFE factorization.

Spectra, expiMap, and SLALOM factor exports were standardized, reduced to 10 or 15 principal components, normalized to unit row length, and evaluated using 5, 10, or 15 cosine neighbors and Leiden resolutions 0.02, 0.1, 0.3, 0.5, and 1.0. The highest cosine silhouette across these 30 configurations was retained without matching the author cluster count. Silhouette calculations used at most 2,000 cells. Thus, the comparator and scKNIFE partition-selection procedures were not identical. Agreement metrics used exact common-barcode intersections of 81,142 cells in C1, 36,264 in C2, and 38,251 in Wu. Runtime summaries came from one recorded 20,000-cell run per method and cohort; the archive did not establish replicate timing uncertainty or a common hardware-controlled experiment.

For Tahoe, MAGIC was run separately within each treatment subset using $t = 3$, the approximate solver, six workers, and `random_state=42`. Scanpy `score_genes` used `ctrl_size=50`, `n_bins=25`, and `use_raw=False`. *PIK3R1* or *TP53* expression was taken from the same MAGIC-imputed treatment representation, whereas scKNIFE activities came from the separately fitted graph model. Pearson and Spearman coefficients were descriptive; no formal test compared the methods' correlation coefficients.

#### Methods S3. Configuration of benchmark latent-variable models

**Spectra**. Spectra was fitted with *est_spectra* to infer cell-type-specific factor matrices[16]. Priors comprised metabolic tasks and biological pathways from Cytopus. Fine-grained Cytopus cell types were mapped to the coarse annotations in each dataset. Gene sets with at most three genes or at least 20% overlap with another set were removed. We set $\lambda = 0.1$, $\delta = 0.001$, and $\rho = 0.001$, left $\kappa$ as `None`, enabled highly variable genes and cell-type-specific factors, and trained for 10,000 epochs. Cell-by-factor embeddings were extracted from `SPECTRA_cell_scores` in the AnnData `.obsm` slot.

**SLALOM**. SLALOM factorized each log-transformed expression matrix with a binary prior compiled from Reactome, Gene Ontology Biological Process, Molecular Function, Cellular Component, and human KEGG gene sets[17]. Gene sets represented by at least 85 genes were retained. Models used annotated factors only (`nHidden=0`, `nHiddenSparse=0`), no pretraining or gene pruning, `minGenes=1` after filtering, and a Gaussian noise model. Other parameters and the 1,000-iteration schedule followed package defaults. The resulting cell-level factor states were used as embeddings.

**expiMap**. expiMap used a conditional variational autoencoder with a linear decoder masked by a gene-program matrix[18]. Reactome, Gene Ontology, and KEGG pathways represented by at least 65 genes in each dataset were retained; genes outside the retained masks were removed, and source prefixes were added before masks were merged. The encoder contained three 256-unit layers. Models used negative-binomial reconstruction, layer normalization, 5% dropout, Adam with learning rate $1 \times 10^{-3}$ and $\varepsilon = 0.01$, group-lasso regularization with $\alpha = 0.7$, KL weight 0.5, and 100 epochs of KL annealing. Training ran for at most 400 epochs with zero weight decay, a 0.9 training fraction, seed 2020, GPU acceleration, and memory-adaptive batches capped at 32,768. Early stopping used patience 50, and the learning rate was reduced tenfold after 13 stagnant epochs. Active cell-resolved pathway factors were extracted with `get_latent(mean=False, only_active=True)`.

#### Methods S4. scKNIFE latent-space clustering and graph-supported cell annotation

The benchmark clustering grid evaluated 10 or 15 principal components and 5, 10, or 15 neighbors. Cohort-specific Leiden resolutions were evaluated with random seed 42. Silhouette coefficients used at most 2,000 cells. Configurations were ranked by silhouette among solutions matching the number of author-defined classes. For Bassez A, an additional unconstrained configuration was selected by maximizing silhouette over the complete grid. This distinguished unconstrained transcriptional heterogeneity from annotation-count-matched benchmarking.

After scKNIFE fitting, the `H_graph` derived from the cell-factor representation was clustered separately from reference annotation. The saved atlas UMAP was computed from the post-fit cell-factor representation $\mathbf{H}^\top$ for visualization and did not define clusters or cell-level SingleR labels.

For benchmark annotation, every available marker-node activity was averaged within each finalized cluster. The marker feature with the highest cluster mean defined the primary label and was propagated to all member cells. Standardization of marker means within clusters provided a quality-control assignment.

For the TNBC cohort, cell-level reference annotation used SingleR v2.4.1[19] and the full Wu breast-cancer atlas, comprising 100,064 reference cells from 26 donors[14]. Reference and query raw counts were independently normalized to 10,000 counts per cell and natural-log1p transformed. The normalization denominator included all 29,733 reference genes and all 16,456 available query genes in their respective matrices; the normalized matrices were then restricted to 16,456 shared genes. Wilcoxon reference-marker detection and within-label reference aggregation with power 0.5 were used for training. The resulting classifier was applied separately to each of the 69,391 query-cell expression profiles, using score quantile 0.8, fine-tuning threshold 0.05, and default SingleR outlier-based pruning. Aggregation applied to the reference; each query cell retained its own prediction.

Major and minor reference labels were predicted independently. Raw predictions, pruned labels, and confidence margins were retained by cell ID; missing pruned labels were recorded as ambiguous. Figure 4 retains the original 12-category annotation alongside independently assigned cell-level full-Wu SingleR annotations. The nine major reference labels were represented as eight broad display groups by combining cancer and normal epithelial predictions into Epithelial. The minor tier retained 29 reference identities after pruning, with 72 ambiguous cells. Cell-level annotations were aligned by cell ID to the saved $H$-derived UMAP for visualization. These labels were not factorization targets. Figure 7 uses 69,319 joined cells, eight broad SingleR groups for the CellChat comparison, and 29 pruned minor identities for the resolved PDGF matrix. Supplemental Table 4C preserves the cell-level crosswalk; the broad and minor tiers are descriptive annotation layers rather than a forced hierarchy.

Canonical-marker expression supported annotation review and was not individually thresholded. Cell states were assigned independently from graph-encoded state-node activities; missing state nodes were excluded rather than imputed. Cluster-by-state scores, cluster mappings, cell-level assignments, and annotated AnnData objects were retained for auditability.

#### Methods S5. Metabolic evidence integration and Human-GEM reaction-rule scoring

MEIA was calculated for 218 scCellFie tasks and 138 analyzable Human-GEM pathways. For task or pathway $e$ in cell $c$, direct entity activity was combined with mean activity over its mapped reactions and genes as

$$Q_e(c) = \big(S_e(c)+\varepsilon\big)\sqrt{\big(\overline{S}_{\mathcal{R}_e}(c)+\varepsilon\big)\big(\overline{S}_{\mathcal{G}_e}(c)+\varepsilon\big)}, \qquad \varepsilon = 10^{-10}. \tag{S1}$$

Task and pathway blocks were normalized separately to sum to one within each cell. Direct scKNIFE entity scores were normalized using the same two blocks before comparison. Raw and normalized values were retained, and a normalized value was interpreted as relative allocation within its block.

Human-GEM identifiers were mapped to HGNC symbols represented in the scKNIFE activity matrix. Gene–protein–reaction (GPR) expressions were parsed independently in every cell. AND relationships used the elementwise minimum activity of required genes, whereas OR relationships used the maximum among alternative genes or isozymes. When multiple Human-GEM reaction records mapped to the same sanitized reaction name, their gene-derived values were averaged. The retained mapping contained 8,043 Human-GEM reaction records corresponding to 7,126 unique scKNIFE reaction names.

For each mapped reaction $r$, the gene-derived score $M_r$ was divided by its reaction-specific 99th percentile and clipped to $[0,1]$:

$$\widetilde{M}_r(c) = \min\left(\frac{M_r(c)}{P_{99}(M_r)}, 1\right). \tag{S2}$$

Reactions were assigned to 138 Human-GEM pathways using model subsystem annotations. The gene-derived pathway score was

$$\text{GPR-GPA}_p^{\text{global}}(c) = \frac{1}{|\mathcal{R}_p|}\sum_{r\in\mathcal{R}_p}\widetilde{M}_r(c), \tag{S3}$$

and the pathway-specific form divided this value by the mean of the other 137 pathway scores:

$$\text{GPR-GPA}_p^{\text{specific}}(c) = \frac{\text{GPR-GPA}_p^{\text{global}}(c)}{\text{mean}_{q\neq p}\big[\text{GPR-GPA}_q^{\text{global}}(c)\big]}. \tag{S4}$$

The focal pathway was excluded from its denominator. The GPR-derived values were compared with direct pathway-node activities and were not multiplied by them.

Technical concordance used exact barcode intersections. MEIA and direct scKNIFE scores shared 69,390 cells and 356 entities; one P3 cell in the 69,391-cell direct atlas was absent from the MEIA intersection. Entity-wise correlations compared cells for each entity. Separately, within-cell correlations compared the 356-entry activity profiles and were summarized by patient medians and interquartile ranges. GPR-GPA-specific and direct scKNIFE values were compared over the 138 pathways. These comparisons evaluated consistency among related transcriptome-derived constructions rather than accuracy against metabolite abundance, reaction rates, or pathway flux.

#### Methods S6. Cell-resolved metabolic activity screening and task prioritization

Cell-level activity comparisons used two-sided Wilcoxon rank-sum or Mann–Whitney $U$ tests within cell type, requiring at least 20 cells per group. The primary task analysis used scKNIFE MEIA scores for 218 tasks; integrated and Human-GEM pathway analyses were sensitivity analyses. Benjamini–Hochberg correction was applied within predefined test families. Because these pooled-cell screens did not account for patient-level replication, they were treated as exploratory. The sulfur-focused study-cohort analysis tested nine scores in five compartments using paired patient-level Wilcoxon tests with correction across 45 comparisons; within-patient time-point comparisons were exploratory and corrected separately.

#### Methods S7. Epithelial node-group construction and activity summaries

The epithelial analysis used 36,963 cells assigned to the broad epithelial group, 50 aligned scKNIFE factors, and nine W-derived node groups (NG1–NG9). Cell-type and cell-state annotation nodes and zero-signal rows were excluded before grouping. For node $i$ and factor group $g$, enrichment was defined as the mean standardized loading inside $g$ minus the mean outside $g$. Each node was assigned once to its strongest positive-enrichment group. Within each group, candidates were ranked using weights $\exp[-(\mathrm{rank}-1)/5]$; ranks through 95% cumulative weight were retained, up to 20 globally unique nodes per group. The 180 selected nodes were used for the display, and functional group names are descriptive summaries rather than independently validated pathways.

Selected node activities were reconstructed from all 50 factors as $\max(\mathbf{H}\mathbf{W}_{\mathrm{node}}, 0)$, transformed with $\ln(1+x)$, and standardized across the epithelial cohort for display. This differs from factor-group usage, which averages the corresponding rows of $\mathbf{H}$. Program-to-factor-group links summarize standardized $\mathbf{W}$ loadings, factor-group-to-epithelial/state links summarize standardized mean $\mathbf{H}$ usage, and epithelial/state-to-patient links represent observed cell counts. Cell-weighted and equal-patient summaries were retained separately. P4 and P1 biopsy/surgery comparisons used cell-type- and RNA-depth-matched cells and are descriptive summaries of distinct sampled populations, not repeated measurements of the same cells or inferential patient-group tests.

#### Methods S8. Patient-resolved treatment-associated metabolic activity summaries

Inferred node and pathway scores were summarized by patient, time point, and cell compartment. Paired biopsy–surgery comparisons used exact two-sided Wilcoxon signed-rank tests across the seven patients with both specimens; patient summaries were the units of inference. Recurrence values from the single contributing patient were descriptive. Benjamini–Hochberg correction was applied separately to the node (1,344 tests), task (1,308 tests), and pathway (828 tests) screening families. Node tests were also adjusted within each of four mapped LC–MS pathways; the focused carnitine analysis tested 60 comparisons with the same correction procedure. Inferred activities are relative transcriptome-derived scores, not measurements of metabolite concentrations or flux.

#### Methods S9. Permutation analysis and CellChat concordance of communication scores

scKNIFE communication scores used evidence weights of 1.0 for KEGG-supported interactions, 0.8 for other documented evidence, and 0.5 when evidence was missing. Cofactor multipliers were 1.5 for agonists, 0.5 for antagonists, 1.3 for co-activating receptors, and 0.7 for co-inhibitory receptors. Pooled tests permuted cell labels 1,000 times while holding specificity weights fixed; one-sided empirical $P$-values were calculated as $(b+1)/(1{,}000+1)$ and adjusted by Benjamini–Hochberg across 252 pathways and 2,261 ligand–receptor keys.

Metastatic-status and sampling-site contrasts were computed within the relevant cell subsets; surgery minus biopsy defined the direction, and recurrence cells were excluded.

CellChat probabilities used saved Seurat-normalized expression, `computeCommunProb` with `truncatedMean` and `trim=0.1`, and at least ten cells per population. scKNIFE and CellChat outputs were harmonized by pathway, ligand–receptor identifier, and broad cell type; overlap was summarized by set counts and Jaccard indices, and matched positive scores were compared on the $\log_{10}$ scale using Pearson and Spearman correlations. Patient-paired biopsy–surgery analyses required at least ten sender and receiver cells at both time points; exact paired sign-flip $P$-values were adjusted across 2,080 pathway/source/target comparisons.

#### Methods S10. Longitudinal Gene Ontology activity analysis in Bassez cohorts

##### GO-node activities, cohorts and metadata

Gene-to-term relationships from the Enrichr Gene Ontology (GO) Biological Process 2021 collection formed a native layer of the scKNIFE knowledge graph. For GO term $g$ and cell $c$, activity was reconstructed from the node-factor loading matrix $\mathbf{W}$ and cell-factor activity matrix $\mathbf{H}$ as

$$A_{gc} = \mathbf{W}_{g,:}\mathbf{H}_{:,c}. \tag{S5}$$

Rows whose names ended in a seven-digit GO accession were extracted from the frozen 22,951-node activity matrices for Bassez cohorts A and B. Both cohorts contained the same ordered universe of 6,036 unique GO Biological Process terms. Columns were aligned to unique cell identifiers in the matched scKNIFE AnnData metadata.

Cells were retained if they belonged to expanded (E) or non-expanded (NE) T-cell clonotype strata, were sampled at pretreatment (Pre) or on-treatment (On), and were assigned to one of six major types: B cell, tumor or epithelial cell, endothelial cell, fibroblast, myeloid cell or T cell. The aligned datasets contained 71,720 cells in Bassez A and 36,527 in Bassez B. E and NE denote clonotype expansion, not clinical response.

##### Patient-level treatment-response interaction analysis

The analyses in Supplementary Figure S3AC–D used patients as the statistical units. For each patient, GO activity was summarized as the median across cells within each cell type and time point, requiring at least 20 cells at both Pre and On. The paired response was calculated as $\log_2$(On/Pre). For each GO term and cell type, limma modeled these patient-level responses by clonotype-expansion group; the NE-minus-E coefficient and its 95% confidence interval summarize the group-by-time interaction. Benjamini–Hochberg correction was applied across all testable GO-term-by-cell-type interactions separately within each cohort. Terms were selected in cohort A by adjusted $q$ value followed by absolute effect, and the same terms were displayed in cohort B. Display redundancy filtering did not change the tested family. These patient-level tests are distinct from the pooled-cell screen below.

##### Pooled-cell differential testing and effect summaries

On was compared with Pre separately within each cohort, expansion group and cell type using Scanpy `rank_genes_groups` with `method=wilcoxon`. The procedure reports a Wilcoxon rank-sum score and a two-sided $P$-value from the normal approximation; cells are the testing units in this pooled-cell screen, and cells are nested within patients. Each contrast required at least 20 cells at both timepoints. All six cell types were tested in Bassez A E, Bassez A NE and Bassez B E. Bassez B NE B cells were excluded because only 15 Pre cells were available.

For each term, the displayed effect was calculated from arithmetic cell-level means as

$$\log_2 \mathrm{FC}_g = \log_2 \left( \frac{\overline{A}_{g,\mathrm{On}}}{\overline{A}_{g,\mathrm{Pre}}} \right). \tag{S6}$$

This value was recorded as missing when the Pre mean was not positive. Cohen's $d$ was calculated as

$$d_g = \frac{\overline{A}_{g,\mathrm{On}} - \overline{A}_{g,\mathrm{Pre}}}{\sqrt{(s^2_{g,\mathrm{On}} + s^2_{g,\mathrm{Pre}})/2}}, \tag{S7}$$

where sample variances used denominator $n-1$. If the denominator of $d_g$ was zero, $d_g$ was set to zero.

Benjamini–Hochberg correction was applied across the complete GO-term-by-cell-type family within each cohort and expansion group, after combining all 256-term processing batches. Bassez A E, Bassez A NE and Bassez B E each comprised 36,216 tests (6,036 terms across six cell types). Bassez B NE comprised 30,180 tests across five cell types. Processing batches controlled memory use and did not define separate correction families.

#### Methods S11. Independent processing and characterization of study-cohort scRNA-seq data

An independent study-cohort scRNA-seq workflow processed 80,217 Cell Ranger-called cells with Scanpy 1.9.6[20]. Quality control excluded cells with fewer than 200 detected genes, more than 20% mitochondrial counts, or precomputed doublet calls, and genes detected in fewer than three cells; 69,391 cells and 16,456 genes remained. Clinical metadata were joined by library–sample identifiers. Library-aware Seurat-v3 selected 3,000 highly variable genes for scVI[21,22] integration; Harmony[23] was assessed as a comparator. Leiden clustering yielded 21 communities at resolution 0.7. Cell identities were assigned using canonical markers, CellTypist[10], PanglaoDB[24], breast/TNBC references, scArches mapping[25], and expression modules, with epithelial-subtype refinement and marker analysis. infercnvpy (version 0.4.5) supported transcriptome-derived copy-number annotations, and Vanderbilt TNBCtype-4 signatures[26,27] supported epithelial-cell subtype scoring. This workflow was independent of scKNIFE; CNV and subtype labels are computational annotations.

SUPPLEMENTAL TABLES 1–7

Supplemental Table 1. scKNIFE factor diagnostics and cross-cohort concordance.

(A) Factor diagnostics for Bassez C1, Bassez C2, Shiao and Wu TNBC, including graph-node and factor counts, effective rank, effective-rank percentage, absolute factor correlations and the condition number of column-standardized W. All four matrices contain the same 22,951 ordered knowledge-graph nodes. Effective rank is $\exp(-\Sigma\, p_i \ln p_i)$, where $p_i = s_i^2/\Sigma\, s_j^2$ and $s_i$ are the singular values of column-standardized W. The effective-rank percentage is effective rank divided by the number of factors. Factor correlations are absolute Pearson correlations across distinct factor pairs. The condition number is $s_{max}/s_{min}$ for standardized W. (B) Six unique pairwise linear centered-kernel alignment (CKA) values. For each comparison, W was column-z-scored, scaled by the square root of its factor count, row-L2-normalized and column-centered before calculating $CKA(X,Y)=||X^TY||^2_F/(||X^TX||_F\ ||Y^TY||_F)$. Comparisons use aligned graph-node rows and do not require matching factors. Bassez C1 and C2 correspond to cohorts A and B, respectively.

Supplemental Table 2. Composition of the benchmarking and in-house knowledge graphs.

Version 1 is the 22,951-node benchmarking graph. Version 2 retains version 1 and adds the Human-GEM component, for a total of 39,247 nodes. The coverage totals of 7,521 and 7,668 represent configured biological-program category memberships, not unique mechanisms or latent factors.

Supplemental Table 3. Benchmarking parameters and biological evidence for Figures 2 and 3.

(A) Final production parameters; (B) retained hyperparameter-tuning summaries; and (C) candidate-level records supporting Figure 2. The 500-iteration value is the configured training budget. (D) Panel-level biological evidence and patient statistics for the baseline and longitudinal candidates in Figure 3e–j, with supplementary heatmap and Gene Ontology context. The baseline associations in rows e and f are ranked exploratory results; the remaining rows describe treatment-stage candidates and supporting literature. Statistical qualifications and evidence categories are provided with the corresponding records; this table is a companion summary, not a new significance analysis.

Supplemental Table 4. Clinical characteristics, cell annotations and expression-based profiles for Figure 4 and Supplementary Figure S4.

(A) Records for seven patients and 15 specimens. (B) Cell-class summaries for the original 12-class annotation, the eight broad Wu-reference SingleR groups in Figure 4c and the 29 pruned minor identities in Figure 4e, including 72 ambiguous minor calls. Patient and specimen summaries reuse cells and are not independent replicates. (C) Cell-level crosswalks between original classes and broad SingleR groups, and between broad and minor calls. Major and minor SingleR tiers were assigned independently; their cross-tabulation is descriptive and does not define a nested taxonomy. (D) The 40 Wu-reference marker positions in Figure 4d, with detection percentages and gene-scaled expression. (E) Canonical-marker expression for the 12 original classes in Supplementary Figure S4c and the 29 minor identities in Supplementary Figure S4g. (F) Transcriptome-derived CNV scores and status by cluster and SingleR annotation. The CNV-low/normal and CNV-high/tumor groups contain 27,421 and 41,970 cells, respectively; cluster-level scores are repeated across cells and are not independent measurements.

Supplemental Table 5. Nine-group epithelial program analysis for Figure 5 and Supplementary Figure S5.

(A) NG1–NG9 memberships of the 50 fitted factors, labeled PG1–PG9 in Figures 5 and S5, with selected-node counts and descriptive functional labels. (B) The 180 globally unique selected nodes, with 20 nodes per group, their source ranks and selection weights. (C) Matched and all-cell P4/P1 node and program estimates. (D) Selected NG1 and NG8 node changes in P4/P1, reported separately for matched and all-cell estimates. (E) Equal-patient reconstructed NG activity by provisional epithelial/state annotation. (F) Patient-level factor-usage summaries linked to the nine-state cell census, comprising 36,963 epithelial cells. (G) Patient and cell-state summaries for the NG7 metabolic tasks displayed as PG7 in Supplementary Figure S5. Activities are model-inferred scores. Biopsy and surgery contain different sampled cells, not longitudinally tracked cells. Patients are identified by public pseudonyms P1–P7.

Supplemental Table 6. Source values for Figure 6 and Supplementary Figure S6.

(A) LC–MS candidate pathways for treatment (21 candidates) and metastatic status (33 candidates), with Human-GEM mappings. Gamma-distribution P values were adjusted by Holm within each candidate family; five treatment pathways and no metastatic-status pathways have adjusted $P<0.05$. For the treatment comparison, 18 source terms map to 14 distinct Human-GEM targets, three terms are unmapped, and the UpSet plot contains 17 harmonized categories. Category overlap indicates annotation coverage, not statistical agreement. (B) Two pooled MEIA/direct-scKNIFE concordance records and 356 entity-level records (218 metabolic tasks and 138 pathways); the pooled task comparison is shown in Figure 6c. Both analyses use related transcriptomic graph inputs. (C) Seven-patient values and statistics for the eight selected Wu-major Myeloid nodes in Figure 6d–k, including a six-patient sensitivity analysis excluding P7. The displayed patient P values are nominal after pathway-led selection; within-pathway and global node-screen BH q values are reported separately. (D) Patient-level values and pathway-screen statistics for the Epithelial fatty-acid activation pathways in Figure 6l–m and the Myeloid peroxisomal carnitine shuttle in Figure 6n, including descriptive P3 recurrence medians shown in Figure 6q–s. The pathway screen includes 828 comparisons; L–N q values are adjusted across this screen. (E) Seven-patient values for Myeloid L-carnitine, L-octanoylcarnitine and octanoyl-CoA; their activity maps are shown in Supplementary Figure S6C–E, while paired values for L-carnitine and octanoyl-CoA are table-only. (F) The 60 tests in the focused carnitine node–cell-type family and their BH q values. (G) Surgery-time cell-type-specificity values retained in the table, paired patient–pathway–cell-type records, and the 72 pathway–cell-type summaries shown in Figure 6t. Figure 6t reports median paired log2(surgery/biopsy) ratios, patient-direction counts, nominal exact two-sided paired Wilcoxon P values and BH q values from the 828-test pathway screen. The surgery-time specificity summary is not included in the current supplementary artwork. (H) The 1,800 pathway–node–cell mapping rows for four LC–MS-linked Human-GEM pathways, representing 1,344 unique scored node–cell paired tests. The table reports shared-reaction graph membership, seven-patient directions, nominal exact paired Wilcoxon P values, within-pathway BH q values and global BH q values calculated once per unique test. Selenocystathionine in Figure 6d is biochemical context and is not included in this direct graph-membership table. (I) Descriptive P3 Myeloid node medians and cell counts for three nodes at biopsy, surgery and recurrence. Figure 6i–k shows biopsy–surgery comparisons only; no recurrence P or q value was calculated. Patients are identified by public pseudonyms P1–P7.

Supplemental Table 7. Filtered communication comparisons and patient-level PDGF analysis for Figure 7.

(A) The filtered CellChat–scKNIFE pathway union: 32 shared, 44 CellChat-only and 33 scKNIFE-only pathways. (B) The harmonized ligand–receptor union: 225 shared, 194 CellChat-only and 260 scKNIFE-only keys. Method-specific significance filters were applied before forming each union; membership describes coverage, not a patient-level test. (C) The 1,917 shared positive source–target/ligand–receptor records. Log10 scores have Pearson r=0.732 and Spearman rank correlation=0.746; the comparison uses the same cohort and curated interaction knowledge in both methods. (D) All source–target entries in the eight-by-eight broad and 29-by-29 minor PDGF matrices. The pooled Endothelial→PVL score is 109.3353; Endothelial *RGS5*→PVL Immature is the highest-scoring positive minor-cell connection (602.385847). (E) Ligand–receptor contributions to the selected major and minor Endothelial→PVL connections. *PDGFB–PDGFRB* contributes 83.493858% and 88.882337% of the respective inferred connection scores; these percentages are not expression fractions. (F) Paired biopsy–surgery values for seven patients for the absolute Endothelial→PVL PDGF score, relative PDGF share, mean endothelial *PDGFB* log1p expression and mean PVL *PDGFRB* log1p expression. Each measure decreases in six patients and increases in P1. At least ten sender and receiver cells were required at both time points; recurrence was excluded. (G) The 2,080 major pathway/source/target biopsy–surgery tests, with exact paired sign-flip P values and BH q values. No test passes q<0.05; the PDGF patient pattern in Figure 7l–n is a descriptive follow-up, not an adjusted differential finding. Patients are identified by public pseudonyms P1–P7.

### SUPPLEMENTAL FIGURE TITLES AND LEGENDS

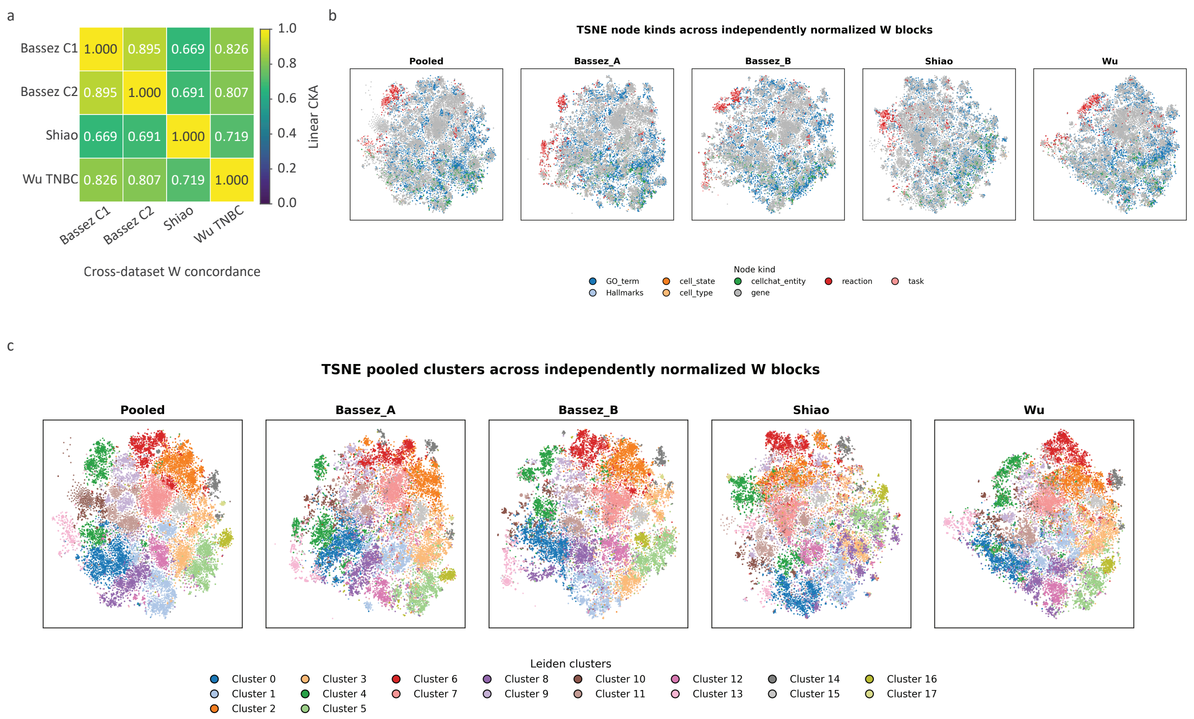


Figure S1

492

**Figure S1. Shared structure in graph-node loading representations.**
(a) Pairwise linear centered kernel alignment (CKA) between the W representations fitted to Bassez C1, Bassez C2, Shiao and Wu TNBC. All comparisons use the same 22,951 graph nodes. (b) Independently fitted t-SNE views of the pooled representation and each cohort-specific W block, colored by graph-node kind. Display order is pooled, Bassez C1, Bassez C2, Shiao and Wu. (c) The same respective coordinates recolored by one 18-community Leiden partition learned from the pooled high-dimensional representation. Points represent knowledge-graph nodes, not sequenced cells. These are descriptive representation summaries without inferential P values or confidence intervals.
Knowledge-node counts describe the shared representation, and CKA coefficients quantify its structural similarity across cohorts.
The cohort-specific W matrices contain 40, 40, 80 and 30 factors, respectively. Each block is column-standardized, divided by the square root of its factor count and row-L2-normalized. Corresponding node rows are concatenated into the pooled representation and row-normalized again. Individual factors are not aligned across cohorts; the joint Leiden labels are transferred to each view.

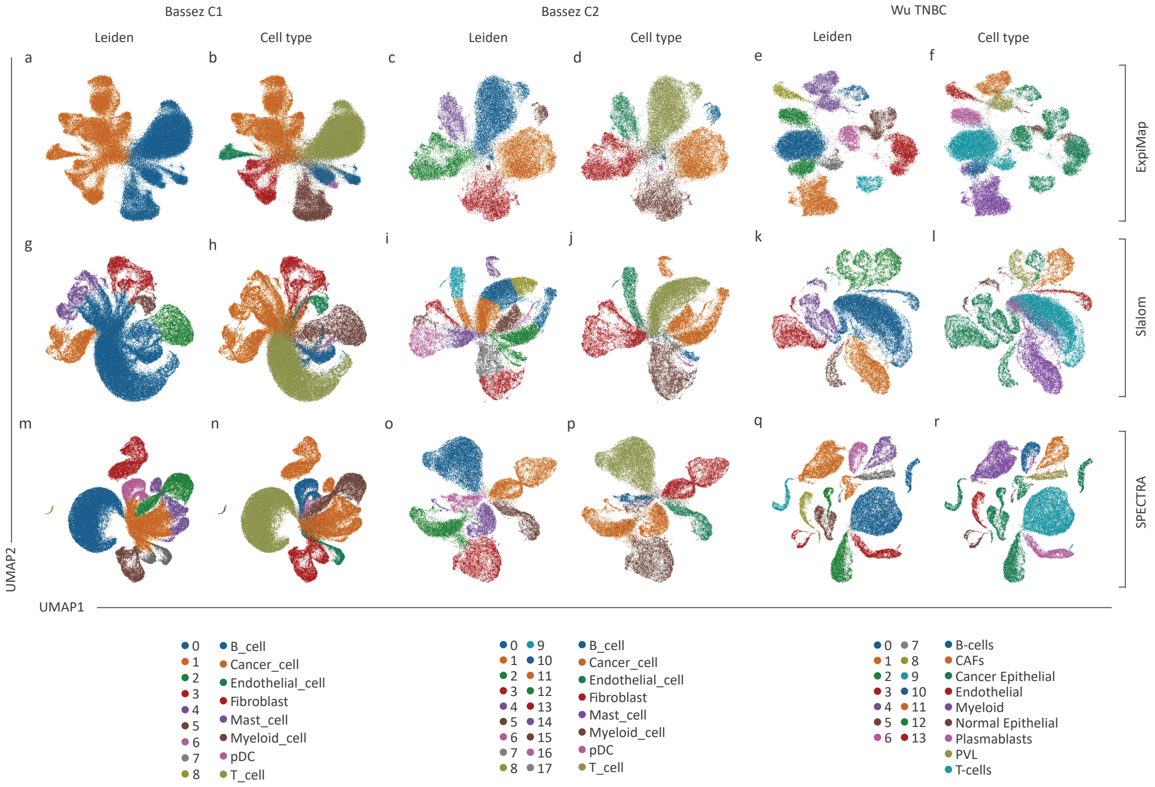


Figure S2

509

**Figure S2. Comparator embeddings document clustering and annotated cell organization across benchmark cohorts.**

Three-by-three grid of frozen comparator embeddings. Rows show expiMap, Slalom and SPECTRA; columns show Bassez cohort 1, Bassez cohort 2 and Wu TNBC. Within each method–dataset combination, the embedding colored by Leiden cluster is shown beside the same coordinates colored by the author-provided cell-type annotation. Each point represents one cell. Embeddings are displayed for qualitative comparison; quantitative agreement metrics are reported in Figure 2l.

**Supplementary Figure 3A · candidate concordance, metabolic tasks and patient-level GO interaction**

Blue: higher R or E · orange: higher NR or NE · outlined points: selected candidates

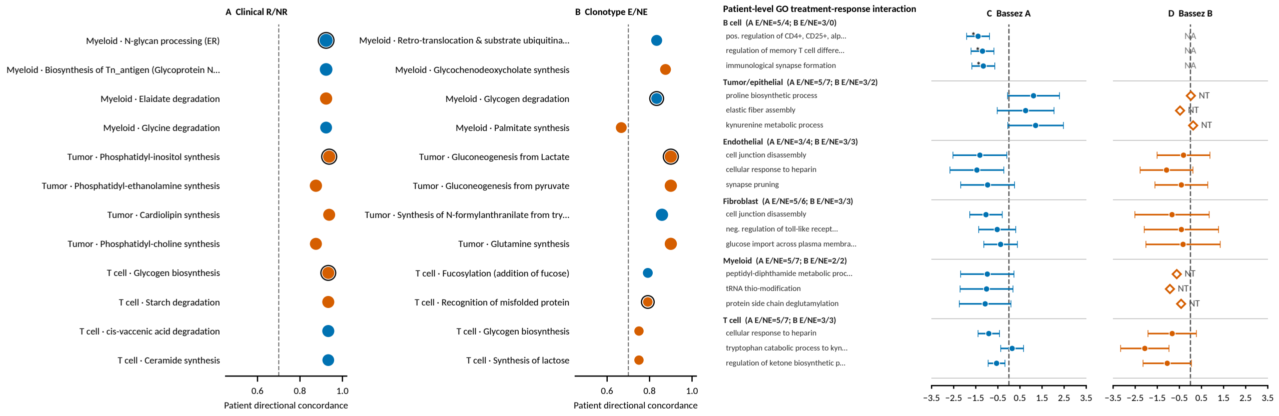


**Pooled-cell metabolic-task log2(On/Pre) · top 10 per group + G/H tasks**

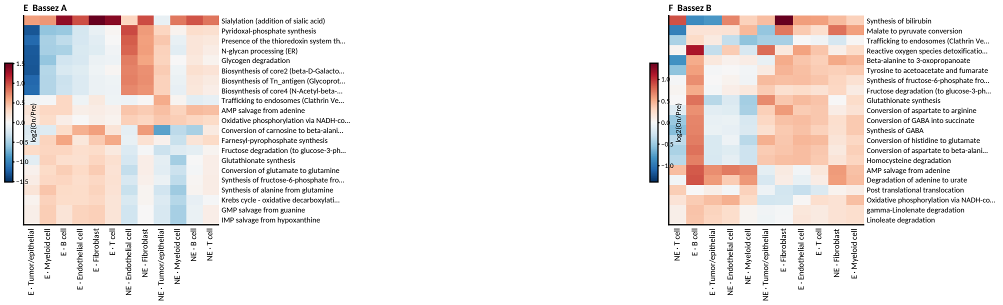


GO: limma estimate and 95% CI · * global BH q<0.05 · diamond: descriptive median (NT) · NA: group absent

**Figure S3A. Candidate concordance and Bassez differential-activity summaries.** (A–B) Patient directional concordance for the leading clinical R/NR baseline candidates across Myeloid, Tumor/epithelial and T-cell compartments (A), and leading Bassez E/NE baseline candidates (B); outlined points mark selected candidates. E/NE denotes T-cell clonotype expansion. (C–D) Patient-level GO treatment-response interaction forest plots for Bassez cohorts A and B, respectively. Within each cell type, GO activity was summarized by the patient median at Pre and On among patients with at least 20 cells at both timepoints, and the paired response was $\log_2$(On/Pre). Points and horizontal bars show the NE-minus-E interaction estimate from a limma model fitted to patient-level responses and its 95% confidence interval. An asterisk indicates global BH $q < 0.05$ across all testable GO-by-cell-type interactions within that cohort. Open diamonds show descriptive median effects when either group has fewer than three paired patients (`NT`); `NA` marks an absent group. Bassez cohort 1 (A) was used to select three nonredundant displayed terms per cell type by global BH q followed by absolute effect, and the same terms were displayed in cohort 2 (B). Display redundancy filtering did not change the tested universe or multiple-testing correction. (E–F) Pooled-cell On-versus-Pre $\log_2$ fold-change heatmaps for metabolic-task activity in Bassez cohorts A and B, respectively. Each heatmap displays the top 10 E and top 10 NE tasks ranked by mean significant $\log_2$ fold change, plus glutathione synthesis and clathrin-dependent endosomal trafficking where available; rows and columns are clustered within the displayed subset. Column labels show the full cell type and E/NE group; `Cancer_cell` is displayed as Tumor/epithelial. The heatmaps are descriptive because cells are nested within patients. GO and metabolic-task values are inferred activities and do not establish pathway flux or causality.

**Supplementary Figure 3B · frozen-embedding task localization**

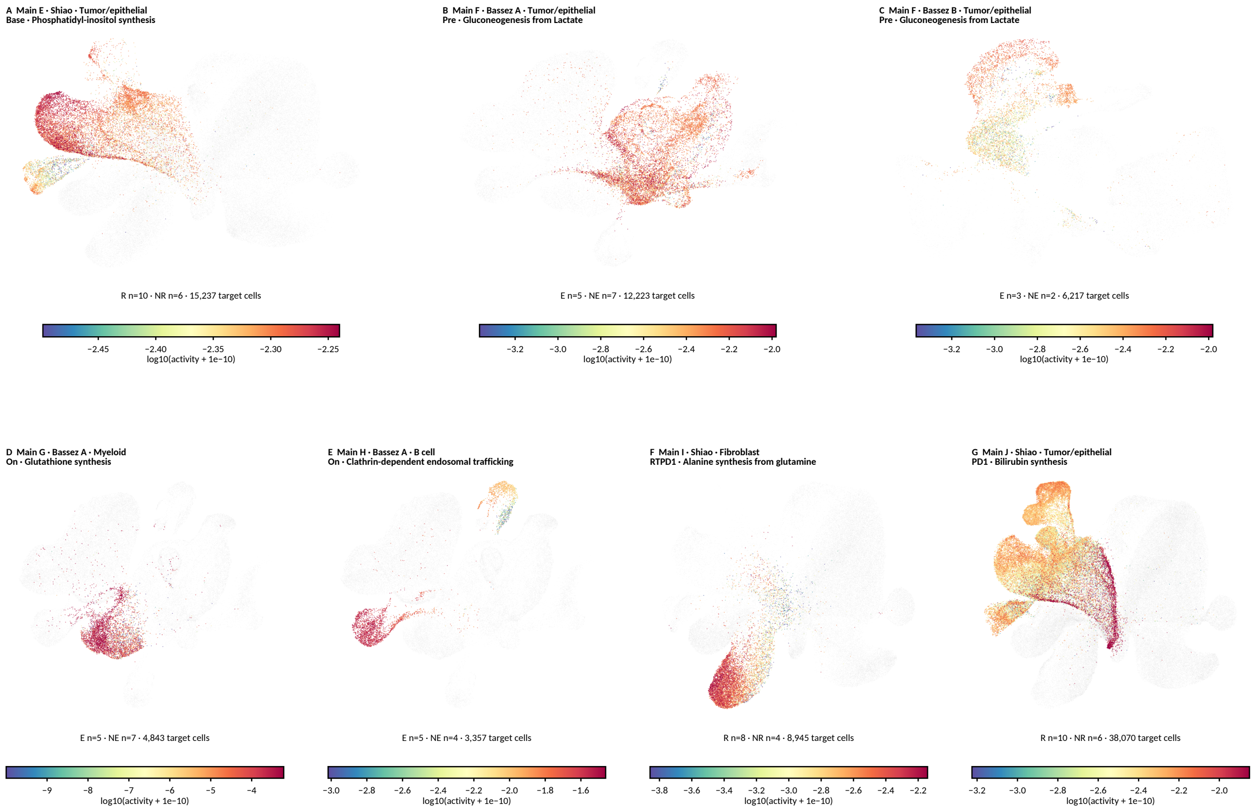

**Figure S3B. Frozen-embedding localization of the activities displayed in main Figure 3e–j.** (A) Shiao baseline Tumor/epithelial phosphatidyl-inositol synthesis corresponding to main e. (B,C) Baseline Tumor/epithelial gluconeogenesis from lactate in Bassez cohorts A and B, corresponding to main f and its supporting cohort comparison. (D) Bassez cohort 1 (A) On-treatment Myeloid glutathione synthesis corresponding to main g. (E) Bassez cohort 1 (A) On-treatment B-cell clathrin-dependent endosomal trafficking corresponding to main h. (F) Shiao RTPD1 Fibroblast alanine synthesis from glutamine corresponding to main i. (G) Shiao PD1 Tumor/epithelial bilirubin synthesis corresponding to main j. Grey points show all endpoint-annotated cells at the displayed timepoint, and coloured target cells show $\log_{10}$-transformed inferred task activity. Outcome groups are pooled in the D–G treated-timepoint maps. Coordinates are frozen cohort-specific UMAP embeddings; colours should be compared only within an association.

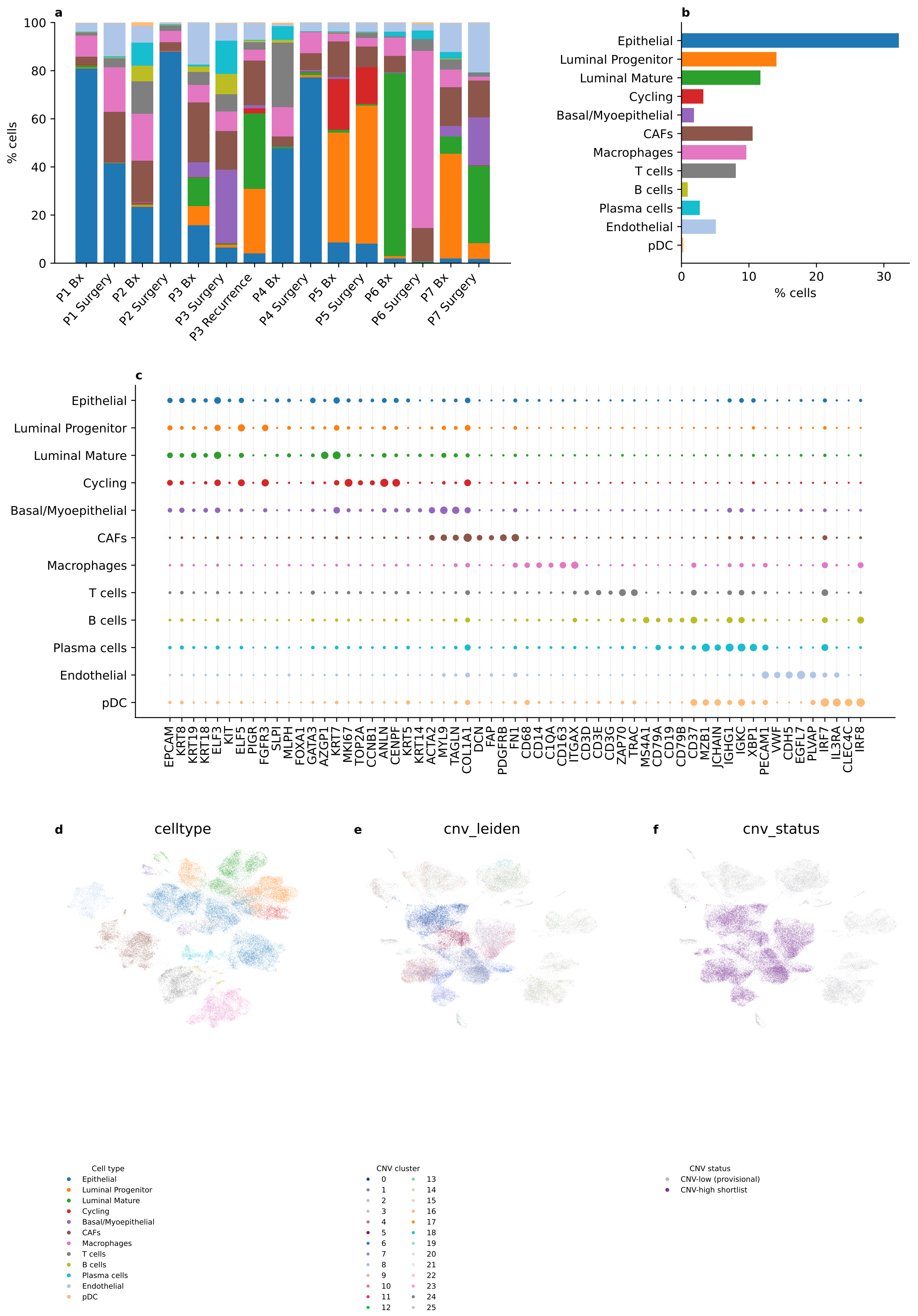
a
100
80
60
40
20
0
% cells
P1 Bx
P1 Surgery
P2 Bx
P2 Surgery
P3 Bx
P3 Surgery
P3 Recurrence
P4 Bx
P4 Surgery
P5 Bx
P5 Surgery
P6 Bx
P6 Surgery
P7 Bx
P7 Surgery
b
Epithelial
Luminal Progenitor
Luminal Mature
Cycling
Basal/Myoepithelial
CAFs
Macrophages
T cells
B cells
Plasma cells
Endothelial
pDC
0
10
20
30
% cells
c
EPCAM KRT8 KRT19 KRT18 ELF3 KIT ELF5 PIGR FGFR3 SLPI MLPH FOXA1 GATA3 AZGP1 KRT7 MKI67 TOP2A CCNB1 ANLN CENPF KRT5 KRT14 ACTA2 MYL9 TAGLN COL1A1 DCN FAP PDGFRB FN1 CD68 CD14 C1QA CD163 ITGAX CD3D CD3E CD3G ZAP70 TRAC MS4A1 CD79A CD19 CD79B CD37 MZB1 JCHAIN IGHG1 IGKC XBP1 PECAM1 VWF CDH5 EGFL7 PLVAP IRF7 IL3RA CLEC4C IRF8
d
celltype
e
cnv_leiden
f
cnv_status
Cell type
Epithelial
Luminal Progenitor
Luminal Mature
Cycling
Basal/Myoepithelial
CAFs
Macrophages
T cells
B cells
Plasma cells
Endothelial
pDC
CNV cluster
0 1 2 3 4 5 6 7 8 9 10 11 12 13 14 15 16 17 18 19 20 21 22 23 24 25
CNV status
CNV-low (provisional)
CNV-high shortlist

555

**g** Reference-derived marker expression across all 29 full-Wu minor identities

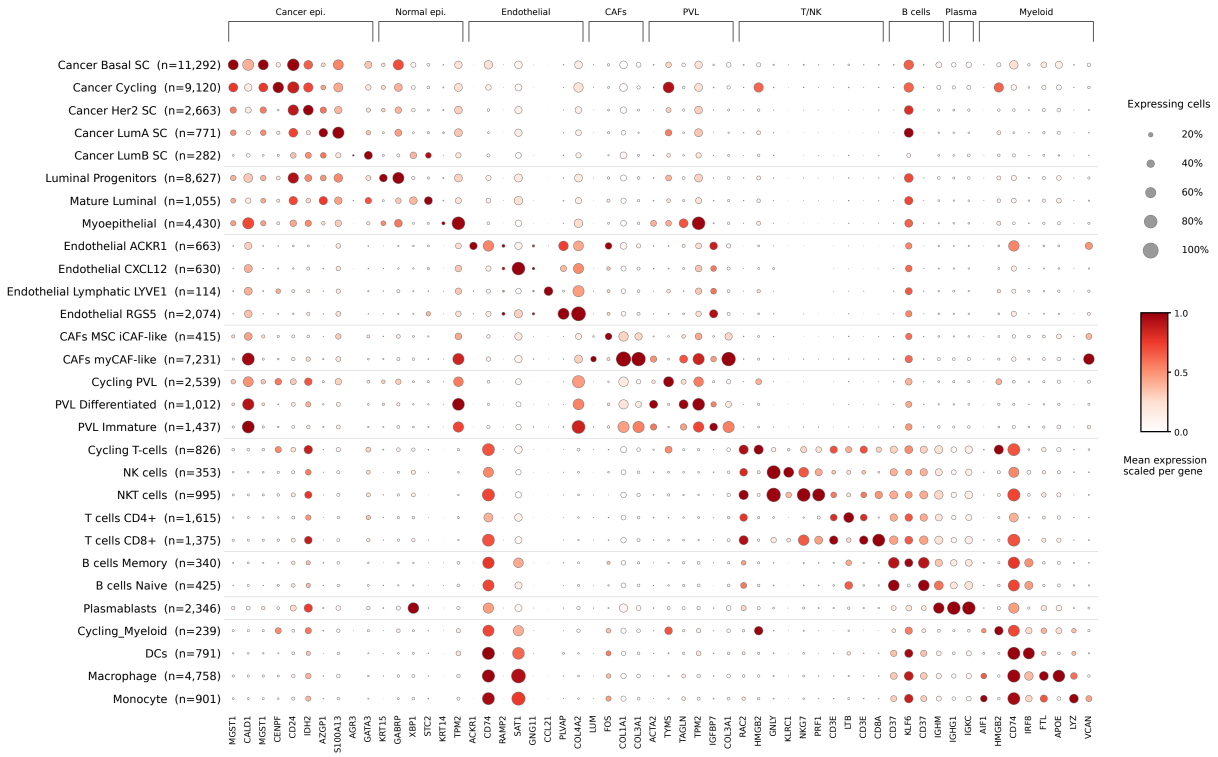


Two saved Wu reference-derived markers per identity; shared genes may repeat. Dot size: cells with expression >0. Color: mean log1p(10k counts/cell), including zeros, scaled per gene.

**Figure S4. Composition, marker and CNV views support annotation of the seven-patient TNBC atlas.**

(A) Stacked percentage composition of the 12 original annotation classes across 15 patient–site or procedure specimens; each bar sums to 100%. (B) Overall percentage composition of the 12 original classes across 69,391 cells. (C) Canonical-marker dot plot showing 59 genes across the 12 original annotation classes. Dot size denotes the fraction of cells expressing each marker, and color identifies the annotated cell type. (D) UMAP of all 69,391 cells colored by the 12 original annotation classes. (E) CNV-QC UMAP colored by 26 expression-derived CNV Leiden clusters. (F) Embedding colored by expression-derived CNV status: CNV-high shortlist ($n = 41,970$) or provisional CNV-low ($n = 27,421$). CNV status is a transcriptome-derived annotation and not an independent DNA-based copy-number measurement.
(G) Full-page white-to-red dot plot across all 29 pruned full-Wu SingleR minor identities, grouped by Wu major lineage. Two markers per identity (58 displayed positions; shared genes may recur) were selected from the saved Wu reference-derived pairwise marker sets: the most recurrent marker and a second marker with the highest within-lineage expression contrast weighted by detection in the query. Dot size indicates the percentage of cells with expression $>$0; color indicates mean log1p-normalized expression, including zeros, min–max scaled separately for each displayed gene. Cell counts appear beside each identity. This panel provides descriptive expression support for the reference predictions.

Figure S5 | epithelial scKNIFE supporting analyses

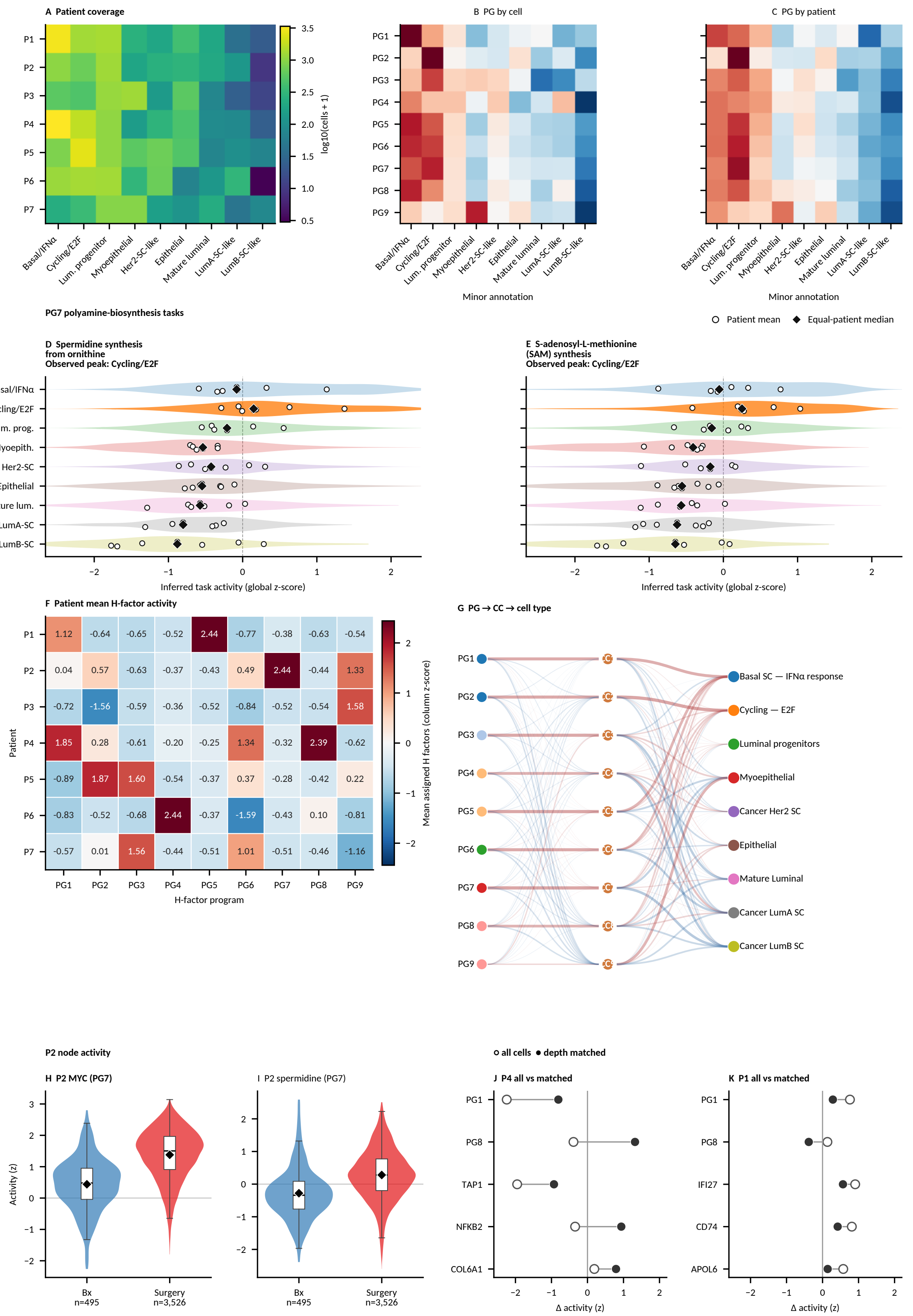

**Figure S5. Supporting analyses of nine epithelial scKNIFE program groups.**
PG1–PG9 correspond to the retained NG1–NG9 analysis identifiers; nine denotes program groups, whereas the epithelial fit contains 50 factors.
(A) Patient coverage by epithelial/state group. (B) Cell-weighted and (C) equal-patient program-by-annotation heatmaps. (D) PG7 spermidine synthesis from ornithine and (E) PG7 S-adenosyl-L-methionine synthesis across epithelial/state groups. Light violins show cell-level inferred activity, hollow circles show patient means, and black diamonds show equal-patient medians. (F) Patient mean assigned H-factor activity, column-z standardized across seven patients. (G) PG-to-component-to-epithelial-label model associations. (H) P2 *MYC* and (I) P2 spermidine-synthesis inferred activities in unmatched biopsy (495 cells) and surgery (3,526 cells) populations. (J) P4 and (K) P1 surgery-minus-biopsy inferred activity contrasting all epithelial cells (open circles) with within-cell-type, RNA-depth-matched cells (filled circles). P4 PG8 changes direction between the estimators. Node activities are log1p-transformed reconstructions standardized across 36,963 epithelial cells; panel F instead summarizes assigned H factors. These descriptive cell-level distributions and activity summaries are not patient-level inferential tests.

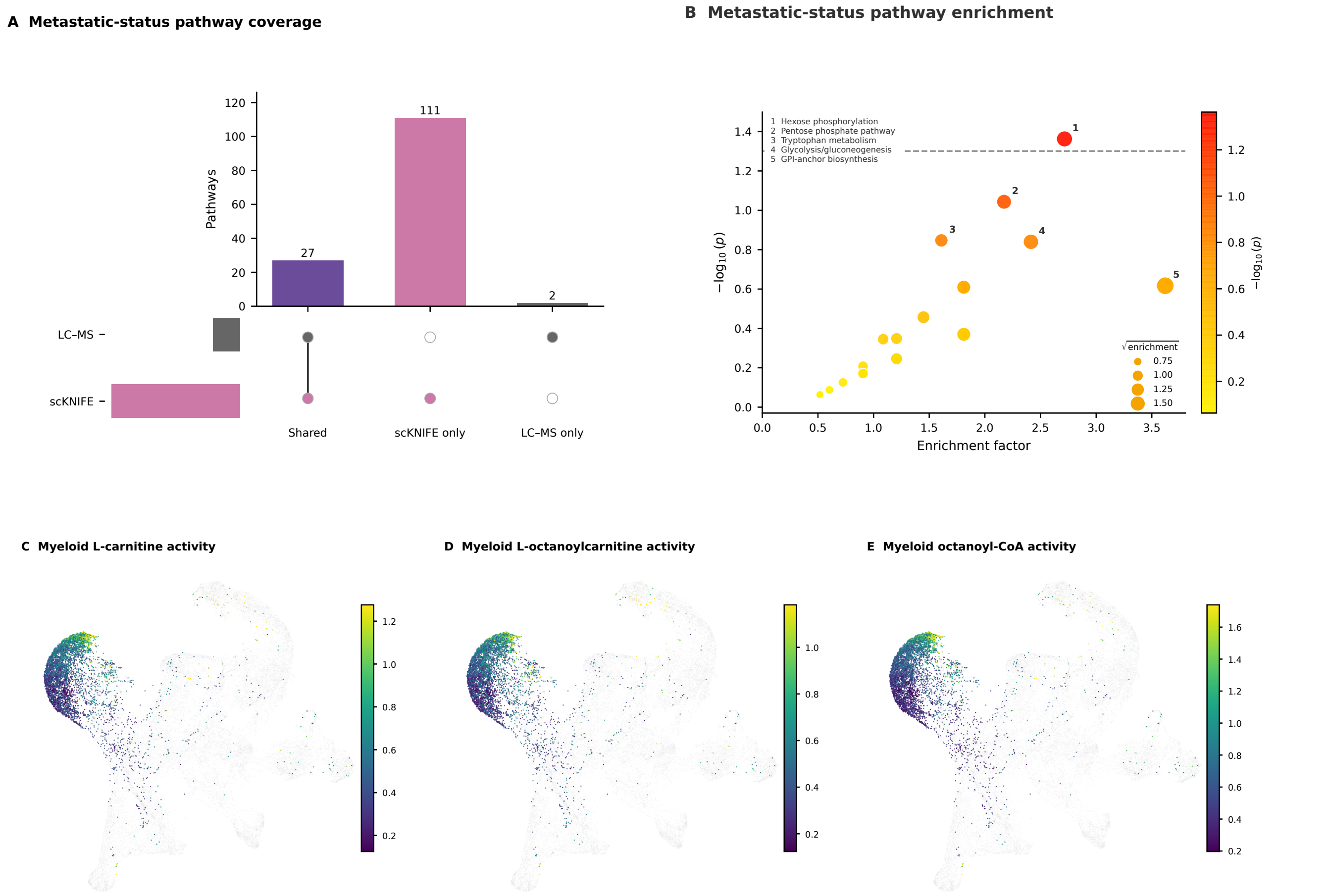
A Metastatic-status pathway coverage
Pathways
120
100
80
60
40
20
0
27
111
2
LC–MS
scKNIFE
Shared
scKNIFE only
LC–MS only
B Metastatic-status pathway enrichment
1 Hexose phosphorylation
2 Pentose phosphate pathway
3 Tryptophan metabolism
4 Glycolysis/gluconeogenesis
5 GPI-anchor biosynthesis
−log10 (p)
Enrichment factor
√enrichment
0.75
1.00
1.25
1.50
C Myeloid L-carnitine activity
D Myeloid L-octanoylcarnitine activity
E Myeloid octanoyl-CoA activity

**Figure S6. Metastatic-status pathway context and Myeloid carnitine activity maps.**
(A) Two-set UpSet plot comparing the 33 LC–MS candidate pathways from the metastatic versus non-metastatic analysis with 138 Human-GEM pathways scored by scKNIFE. Of the 33 LC–MS candidates, 31 map to 27 distinct Human-GEM/scKNIFE pathways; two have no mapping. The display therefore contains 27 shared pathways, 111 scKNIFE-only pathways, and two unmapped LC–MS-only pathways. The LC–MS set size is 29 harmonized terms, reflecting the many-to-one mapping. Membership denotes analytical coverage, not a significant metastatic-status association or cross-modal direction agreement.
(B) Existing LC–MS metastatic versus non-metastatic candidate pathway enrichment scatter for the same 33 LC–MS pathways. The horizontal line marks nominal $p = 0.05$; point color reflects $-\log_{10}(p)$, and point size reflects the square root of enrichment. Numbers label the five leading pathways. None passes Holm-adjusted Gamma $P < 0.05$. The enrichment is exploratory and does not establish pathway direction or cell-type source.
(C–E) The saved Figure 4 H-derived whole-cell UMAP coordinates with all 6,014 biopsy and surgery Wu SingleR major Myeloid cells overlaid by inferred scKNIFE activity scores for L-carnitine (C), L-octanoylcarnitine (D), and octanoyl-CoA (E). The grey underlay includes all 69,391 atlas cells, aligned to Figure 4 by verified cell-ID order. Each activity map has its own 2nd–98th percentile color scale; colors should be compared within a map. Main Figure 6o maps L-octanoylcarnitine, and Figure 6p summarizes its paired patient medians. Paired L-carnitine and octanoyl-CoA values are retained in Supplemental Table 6E.
LC–MS, liquid chromatography–mass spectrometry.
